# HPVarcall: Calling lineages and sublineages for partial DNA sequences of human papillomavirus

**DOI:** 10.64898/2025.12.31.697187

**Authors:** Alexandre Lomsadze, Mark Borodovsky

## Abstract

We describe a computational method, HPVarcall, that assigns DNA sequences of a human papillomavirus (HPV) variant of known type to lineages and sublineages. The algorithm relies on statistical models - positional frequency profiles - trained on multiple alignments of HPV genomic sequences that are known to belong to specific sublineages of a given HPV type.

The workflow begins with multiple alignment of all available sequences for the HPV type, followed by construction of a phylogenetic tree and identification of branches containing sublineage-specific reference sequences. In the prediction phase, sublineage-specific statistical models are used to compute the posterior probabilities for each sublineage given a query sequence. The query classifies to belong to the sublineage with the highest posterior probability. Accuracy assessments performed for the nine HPV types included in the Gardasil 9 vaccine demonstrated a low error rate in assigning HPV genomic fragments of at least 1000 nucleotides to their correct sublineages and even higher accuracy for longer sequence fragments.

## Introduction

The evolutionary history of papillomaviruses spans more than 300 million years (Herbst et al. 2009). The papillomavirus genomes evolve slowly because their double-stranded DNA is replicated by the host’s highly accurate DNA polymerase. Nonetheless, in the relatively young clade of the human papillomaviruses (HPVs) the interplay of mutation and selection along with human population migrations has driven diversification into more than 400 HPV types (Van Doorslaer et al. 2017).

Evidence for the intratype variation of HPV types was found as early as 1981 (de Villiers, Gissmann, and zur Hausen 1981). Advances in sequencing technologies over the past two decades have enabled the large-scale collection of HPV genomic sequences, both complete and partial. Analysis of sequence similarities between HPV types revealed clustering patterns that correspond to lineages and sublineages.

In the current classification system, genomic sequences from different lineages within the same type typically show 1.0–10.0% divergence in average nucleotide identity (ANI) (Chen et al. 2011; Burk, Harari, and Chen 2013). Sequences from different sublineages within the same lineage typically exhibit 0.5–1.0% divergence.

Clinical studies have established that HPV infections carry varying risks of cervical cancers and other malignancies depending on the HPV types (Schiffman et al. 2016). HPV-16, the most carcinogenic HPV type, is associated with approximately 50% of cervical cancers (Mirabello et al. 2016). Further research has demonstrated that cancer risks differ even among lineages and sublineages of the same HPV type. For example, within HPV-16, lineage D carries the highest oncogenic risk, followed by lineage A, with sublineages D2/D3 and A4 considered the most hazardous (Mirabello et al. 2016). Variations in oncogenic potential were also observed among lineages and sublineages of other high-risk HPV types.

These clinically important differences highlight the need for computational tools capable of accurately classifying HPV genomic fragments (often incomplete) obtained from clinical or epidemiological samples. Current methods for variant calling typically rely on phylogenetic tree reconstruction to assign new sequences to specific branches (Burk, Harari, and Chen 2013; Cullen et al. 2015). Yet another approach based on lineage and sublineage specific SNPs (DePristo et al. 2011; Ou et al. 2021; Asensio-Puig, Alemany, and Pavon 2022) was proposed. Application of that approach was limited to discrimination between lineages of HPV-16.

Here, we present a new computational method, HPVarcall, for assigning incomplete HPV sequences to their corresponding lineages and sublineages. The method assumes that the HPV type is known beforehand, e.g. from a prior application of an HPV genotyping tool. The proposed variant calling algorithm uses positional frequency profiles - statistical models trained on multiple alignments of sequences belonging to the same sublineage. Note that the profiles automatically capture information on lineage-specific and sublineage-specific SNPs, reflecting genomic positions with the greatest divergence between sublineages.

Unlike other methods, the new algorithm does not require the input sequence to originate from a specific genomic region like E6 gene of the HPV genome (Cornet et al. 2012). Once the profiles are trained, the computational complexity of the prediction step (taxonomic classification) is comparable to one of the pairwise alignment-based methods. We assessed accuracy using taxonomically characterized HPV fragments and observed good performance for fragments as short as 1,000 nucleotides, with even higher accuracy for fragments of 2,000 nucleotides.

Overall, the proposed method facilitates the detection of high-risk HPV variants by automating HPV taxonomic diagnostics and accelerating accurate variant calling.

## Materials

The Gardasil 9 vaccine targets nine types of the HPV virus: HPV-6, -11, -16, -18, -31, -33, -45, - 52, and -58. For each of the types substantial numbers of complete and partial HPV genomic are available in GenBank with many of them classified in literature into lineages and sublineages. Our work concentrated on these nine HPV types.

In May 2025, 26,422 genomic sequences of HPV variants identified by the type-specific taxonomic IDs were downloaded from GenBank: HPV-6 (10600, 31552, 37122, and 66311), HPV-11 (10580), HPV-16 (333760), HPV-18 (333761), HPV-31 (10585), HPV-33 (10586), HPV-45 (10593), HPV-52 (10618), and HPV-58 (10598).

Algorithm training requires nearly complete genomes. We excluded sequences from GenBank’s patent section, synthetic and environmental (metagenomic) sources, as well as sequences shorter than 7,000 nt, giving us 7,864 sequences (Table 1). Only one representative of genomes with identical sequences was retained. Additional filtering steps for selection of training sequences made upon completing multiple alignments and reconstructions of the evolutionary trees are described in Methods section (see Suppl. Table 1 for the details).

**Table 1.**
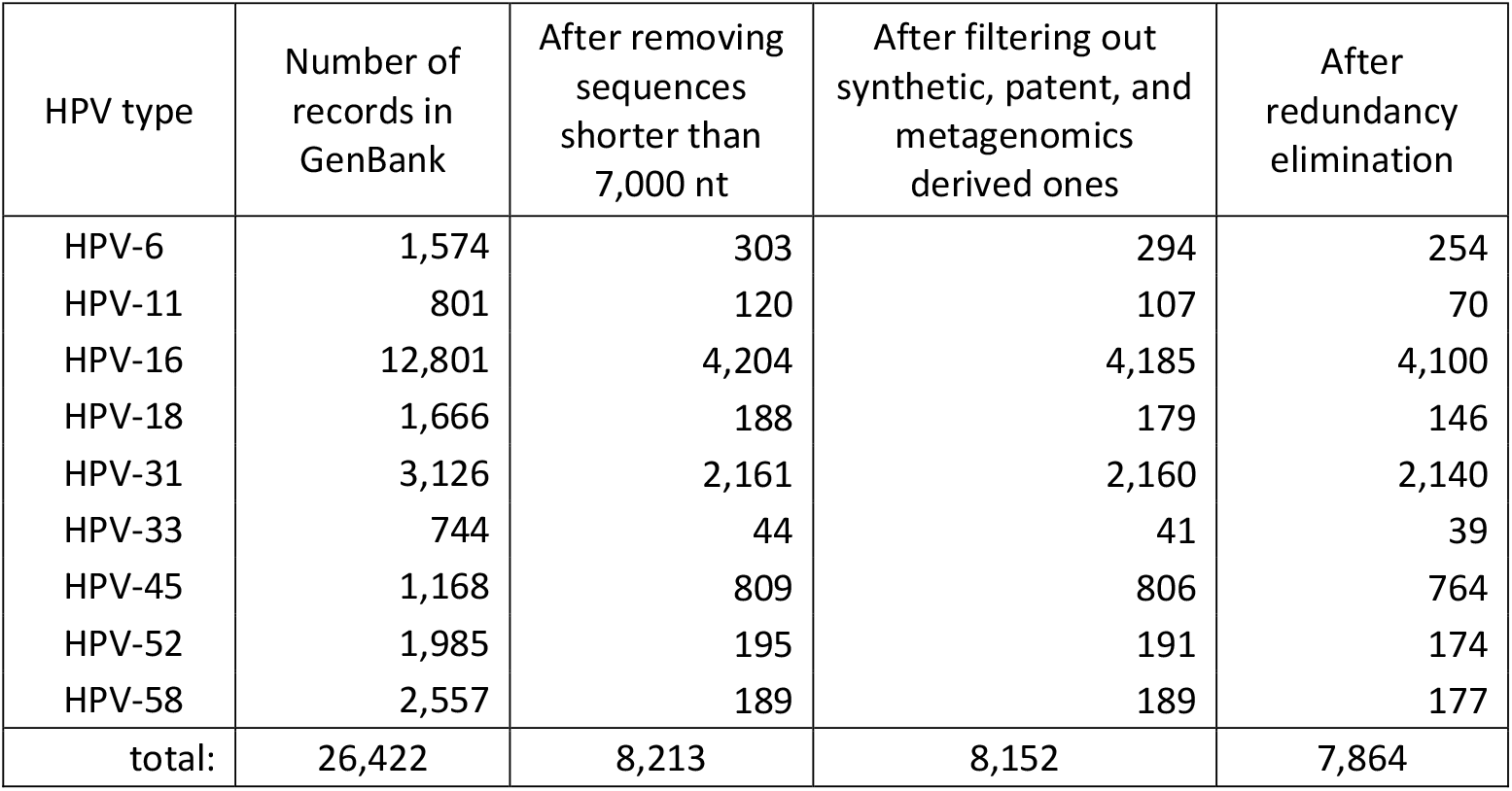
Numbers of selected genomes of each HPV type prior to MSAs construction. The GenBank IDs of the chosen records are listed in Suppl. Table 1.

| HPV type | Number of records in GenBank | After removing sequences shorter than 7,000 nt | After filtering out synthetic, patent, and metagenomics derived ones | After redundancy elimination |
| --- | --- | --- | --- | --- |
| HPV-6 | 1,574 | 303 | 294 | 254 |
| HPV-11 | 801 | 120 | 107 | 70 |
| HPV-16 | 12,801 | 4,204 | 4,185 | 4,100 |
| HPV-18 | 1,666 | 188 | 179 | 146 |
| HPV-31 | 3,126 | 2,161 | 2,160 | 2,140 |
| HPV-33 | 744 | 44 | 41 | 39 |
| HPV-45 | 1,168 | 809 | 806 | 764 |
| HPV-52 | 1,985 | 195 | 191 | 174 |
| HPV-58 | 2,557 | 189 | 189 | 177 |
| total: | 26,422 | 8,213 | 8,152 | 7,864 |

Nine reference genomes of the nine HPV types, along with their gene annotations, were obtained from the PaVE database (Dommer et al. 2025). We should mention that PaVE carries updated reference sequences of HPV types 6, 16, and 18 compared to the original sequences available in GenBank (Van Doorslaer et al. 2013).

Nomenclature of lineages and sublineages of each of the nine HPV types was also acquired from PaVE (Table 2). The sequences of the reference genomes of all the sublineages (with the GenBank IDs, if available, were obtained from PaVE as well (Suppl. Table 2).

**Table 2.** Numbers of lineages and sublineages in the nine HPV types.

| HPV type | Number of lineages | Total number of sublineages | Number of sublineages in each lineage |  |  |  |  |
| --- | --- | --- | --- | --- | --- | --- | --- |
|  |  |  | A | B | C | D | E |
| HPV-6 | 2 | 6 | 1 | 5 |  |  |  |
| HPV-11 | 2 | 5 | 4 | 1 |  |  |  |
| HPV-16 | 4 | 16 | 4 | 4 | 4 | 4 |  |
| HPV-18 | 3 | 9 | 5 | 3 | 1 |  |  |
| HPV-31 | 3 | 8 | 2 | 2 | 4 |  |  |
| HPV-33 | 3 | 5 | 3 | 1 | 1 |  |  |
| HPV-45 | 2 | 8 | 3 | 3 | 2 |  |  |
| HPV-52 | 5 | 9 | 2 | 3 | 2 | 1 | 1 |
| HPV-58 | 4 | 8 | 3 | 2 | 1 | 2 |  |

Reference genomes of seven sublineages currently lack GenBank IDs: A3, A4, and B1 in HPV-11; C4 in HPV-31; and B3, C1, and C2 in HPV-45. We obtained reference genomic sequences of three sublineages for HPV-11 from the authors of the variant classification publications. We retrieved the GenBank IDs of two reference genomes of HPV-45 sublineages using their isolate IDs. Unfortunately, we were unable to locate the GenBank IDs for the reference genomes of C1 sublineage of HPV-31 and B3 sublineage of HPV-45. Therefore, these two sublineages were excluded from our analysis.

We found that the reference genomes of sublineage C3 of HPV-16 (KU053921) and sublineage D1 of HPV-52 (HQ537748) contained indels in protein-coding regions. We replaced these genomes with the closest non-problematic HPV genomes from the same sublineages.

## Methods

### Overall Approach and Role of Multiple Sequence Alignment (MSA)

Finding an optimal multiple sequence alignment under given scoring conditions is a classical problem in bioinformatics. Because the computational complexity of MSA is NP-hard, no algorithm can guarantee an optimal solution for large sets of long sequences. In our study, we construct MSAs for all selected genomic variants of the same HPV type. These alignments serve as a basis for phylogenetic tree construction and subsequent clustering of variants into lineages and sublineages. The MSAs of the sublineage sequences then serve as the input for building probabilistic sublineage-specific profiles.

As noted in the introduction, genomic variants of the same HPV type differ only at a limited number of positions. This high overall similarity greatly simplifies the MSA construction. Note that beyond enabling variant classification, the MSA construction also allows detection of inconsistent genome assemblies, identification of sequencing errors and in some cases correction of these errors.

### The MSA Construction Algorithm

An HPV genome is a double-stranded circular DNA molecule with all protein-coding genes located in the same strand. For analysis, each genome sequence was linearized with the coordinate origine placed at the start of gene E6.

For closely related HPV genomes, we developed a task specific algorithm, fast-MSA which operates in two stages: i/ multiple alignment of all protein coding regions and ii/ multiple alignment of all intergenic regions.

### Multiple Alignment of Protein-Coding Regions

To extract coding sequences, we used gene annotations from the designated reference genome of each HPV type. As a rule, the reference genome of an HPV type corresponds to the reference genome of A1 sublineage of that type (Dommer et al. 2025).

Each genome variant was aligned to the coding regions of its reference genome using BLASTN alignment algorithm. Owing to the high similarity of HPV variants, the most homologous pairs of coding regions would be aligned without gaps from end to end. These ungapped alignments were used to construct the primary MSA blocks for each gene.

Some query sequences could be aligned to reference coding regions with indels. We observed that most indels were deletions, forming gaps in MSA. Single nucleotide deletions within coding regions were interpreted as sequencing errors; the affected positions were replaced with N to preserve the coding frame. Deletions of lengths divisible by three were retained as true biological deletions and represented as gaps in the alignment. Deletions of non-triplet length were interpreted as evidence of assembly artifacts, and such genomes were removed from the dataset (Suppl. Table 3).

### Multiple Alignment of Intergenic Regions

Intergenic regions were inferred from the annotated boundaries of protein-coding genes. All intergenic regions ≥ 21 nt from each genome variant were aligned to the corresponding intergenic region of the HPV type reference genome. As with coding regions, ungapped alignments were first used to construct alignment blocks. Sequences containing deletions relative to the ungapped block were retained, whereas sequences containing insertions were removed.

We observed that some intergenic regions contained low complexity regions that tended to introduce numerous alignment gaps. For these cases, additional filtering steps were applied. The alignment was initiated from the left boundary of the intergenic region and extended until a mismatch/indel threshold was reached. The same procedure was then performed from the right boundary, aligning in the opposite direction. The central segment where both directional alignments exceeded the mismatch threshold was removed, and minimal-length alignment blocks were constructed from the retained flanking segments.

For intergenic regions < 21nt the following was observed. For HPV-6, -11, -18, -52 and -58 all short intergenic regions exhibited uniform length across all variants. For HPV-16, -31, -33, -45, a small number of outliers with length differing from the modal value were detected and removed. This filtering enabled the construction of alignment blocks for the short intergenic regions.

### Phylogenetic Tree Construction and Identification of Sublineage-Specific Branches

All alignment blocks (coding and intergenic) created for variants of a given HPV type were concatenated to produce the whole-genome MSA.

These MSAs were used as input to RAxML (Stamatakis 2014) for maximum-likelihood (ML) phylogenetic tree construction. To identify groups of leaves (variants) belonging to lineages and sublineages, we performed a bottom-up traversal of each phylogenetic tree, beginning at leaves corresponding to established sublineage reference sequences.

The ‘ancestor’ of two branches (sublineages) was assigned to the internal node where the paths ascending from the reference sublineage leaves coalesced. All the paths, initiated from sublineage-specific reference genomes would eventually merge at a node representing the ‘ancestor’ of the whole lineage.

Technically, this strategy defines a semi-supervised clustering method, using known sublineage reference sequences to guide the selection of non-overlapping sets of leaves. The resulting clusters correspond to HPV lineages and sublineages which largely conform to classifications reported in prior studies as we have observed in the results of practical implementation of this approach (Suppl. Table 4).

Note that on one occasion (HPV-45), reference sequences in PaVE were ambiguously defined; in this case, substitute reference variants were selected.

As was mentioned, the filtering of input genomes continued throughout the MSA and phylogenetic analysis stages. A substantial number of genomes were removed during MSA construction, primarily due to presence of extended runs of unresolved bases (N). After tree construction and sublineage labeling, few additional genomes were excluded when their placement in the tree did not conform to the PaVE-established lineage and sublineage structure - that is, when the corresponding leaves fell outside the appropriate clusters defined by the reference variants (see Suppl. Table 3).

### Construction of Sublineage-Specific Probabilistic Profiles

A *probabilistic profile* is defined by positional nucleotide probabilities computed across an MSA. For a sublineage-specific profile, these probabilities are estimated directly from the MSA of the variants assigned to that sublineage. Let *t*_*k*_ denote sublineage *k* within lineage *t*. For each alignment position *i* and nucleotide *x* ∈ {*A, C, G, T*}, let *Count*_*i*_*(x)* be the number of occurrences of nucleotide *x* at position *i* in the sublineage-specific MSA. The positional probability of nucleotide *x* at position *i* is then estimated as:

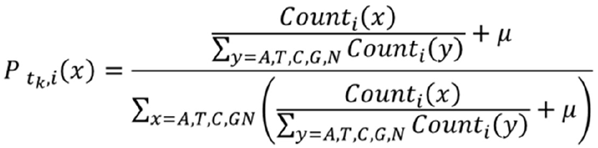

Here µ is a regularization parameter, introduced to regularize the computations for small number of sequences in a sublineage-specific MSA, the µ value was choosen to be 0.005. The whole workflow of the probabilistic profile derivation is illustrated in Fig. 1.

**Figure 1.**
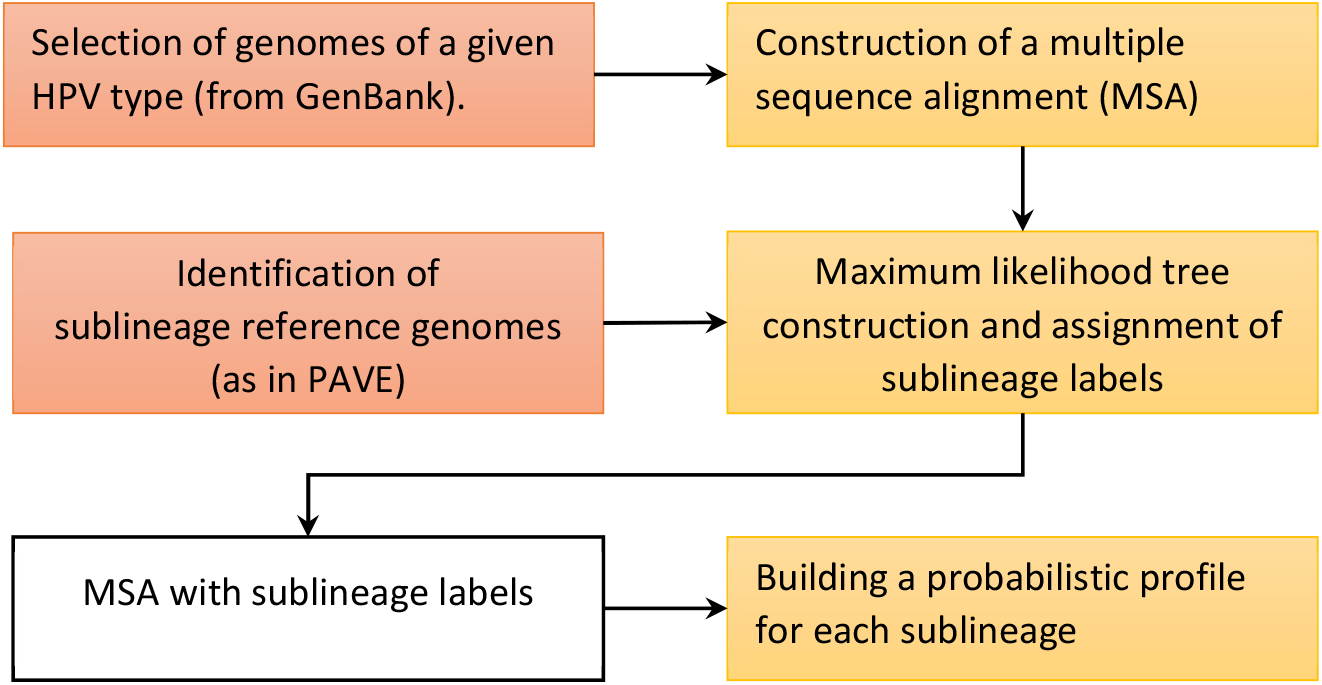
Flowchart of the probabilistic profile construction (model training)

### HPV Variant Calling for a DNA Sequence Extracted from a Sample

During the pre-processing step, the HPV type of the query sequence *s* (genomic fragment) must first be identified by an HPV genotyping tool. The strand orientation of the query sequence is then determined by aligning both the sequence and its reverse complement to the reference genome of the identified HPV type. Once the correct orientation is established, the query sequence is aligned to each sublineage-specific probabilistic profile constructed for that HPV type.

These sequence-to-profile alignments select sections of the profiles that correspond to the query sequence. The posterior probability of each sublineage when observing sequence *s* is then computed using the Bayesian formula:

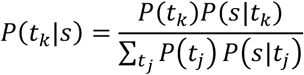

Here *t*_*k*_ and *t*_*j*_ denote candidate sublineages, e.g. *A*_l_ *or B*_2_. The likelihood term, 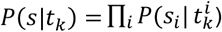 defines the probability of observing sequence *s* in the class defined by the model (profile) of sublineage *t*_*k*_. *P*(*t*_*k*_|*s*) is the posterior probability of sublineage *t*_*k*_ given sequence *s*. The sublineage with the largest posterior probability *P*(*t*_*k*_|*s*) is assigned to the query sequence (Fig. 2).

**Figure 2.**
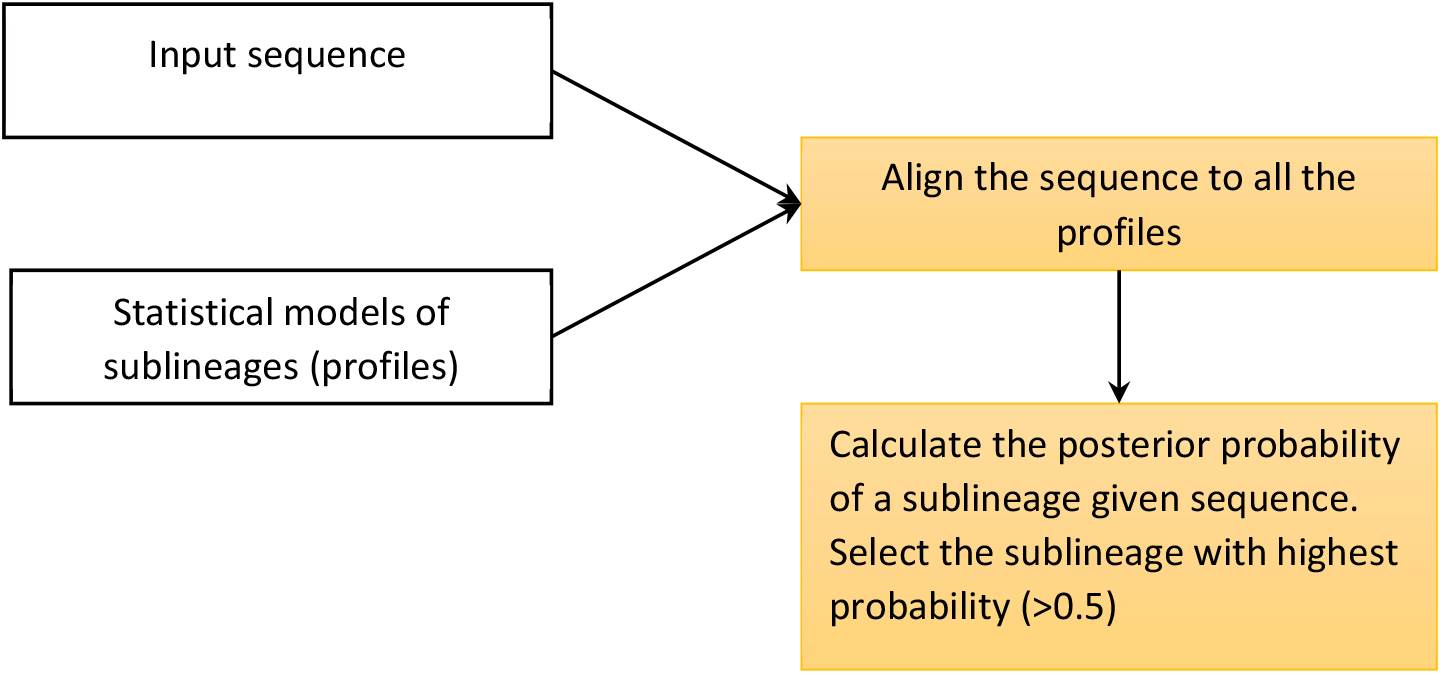
Flowchart of the HPV variant classification (sublineage assignment)

### Assessment of the Algorithm Performance

Incomplete HPV sequences - with length less than 7,000 nt - made approximately 70% of all HPV entries in GenBank in May 2025 (Table 1). Evaluating performance of the proposed method on incomplete genomic fragments is of substantial practical importance.

We assessed the performance of the algorithm on the test sets derived from the sequences of the nine HPV genomes. The lineages, and sublineages of these sequences were assigned according to positions of these genomes in the maximum-likelihood phylogenetic trees. For each HPV type, we generated two test sets of short sequences: 4,000 fragments of length 1,000 nt and 4,000 fragments of length 2,000 nt. These fragments were sampled at random from HPV genomes representing different sublineages. To confirm that 4,000 sequences were sufficient for reliable performance estimates, we also made larger test sets of 50,000 sequences.

## Results and Discussion

### Construction of Phylogenetic Trees and Final Selection of the Training Set

Phylogenetic trees for the nine HPV types and selected 3,306 HPV genomic variants (Suppl. Table 3) were constructed using the fast-MSA algorithm for multiple sequence alignment. Next, the ML trees were built using RAxML. As described in Methods the process of the training set selection was continued upon MSAs building for each type as well as upon sublineages assignments based on trees topologies (Suppl. Table 3). The final versions of the ML trees for the total of 3,272 variants visualized using FigTree (Rambaut 2025) are shown in Fig. 3 and in Suppl. Fig. 1.

**Figure 3.**
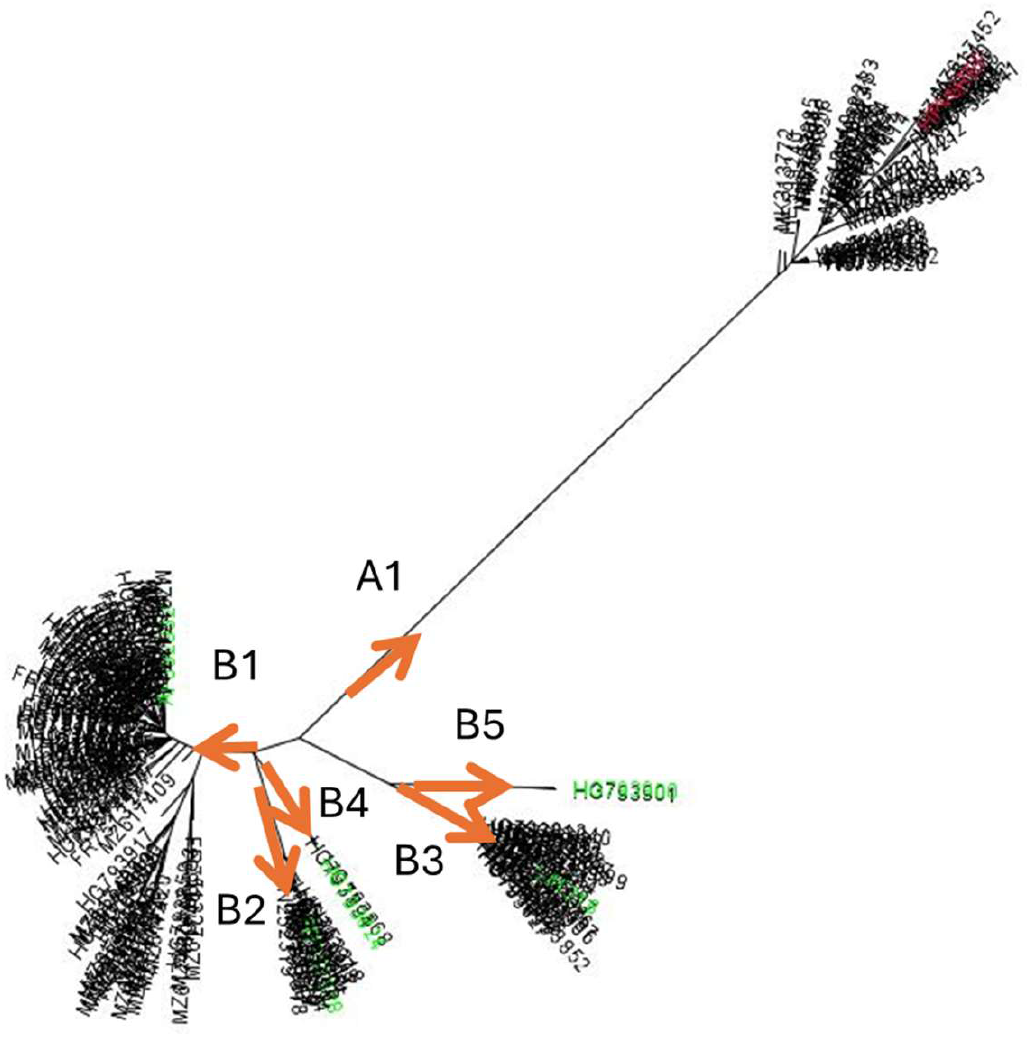
Maximum-likelihood phylogenetic tree constructed for HPV-6 genomic variants, with branches corresponding to established sublineages. Names of reference genomes are color-coded: red for lineage A and green for lineage B.

To ensure that fast-MSA did not introduce systematic bias, we compared the positions of sublineage reference genomes in the constructed ML trees (Suppl. Fig. 1) with those in previously published studies (Burk, Harari, and Chen 2013). No significant discrepancies were observed.

For HPV-6 and HPV-18, we conducted even more detailed comparison. We confirmed that the genomes whose taxonomic affiliation was controlled by manual curation shared the same sublineage assignments as in our work.

In total, 3,272 HPV genomes were classified into lineages and sublineages across the nine HPV types analyzed (Suppl. Table 4.)

### Computation of ANI Divergence for HPV Lineages and Sublineages

Average ANI divergence, computed as 100% - ANI, was determined for reference sequences of all lineages and sublineages across the nine HPV types (Suppl. Table 5). In most cases, ANI divergence between lineages of a given HPV type exceeded 1%, whereas sublineages belonging to the same lineage were separated by smaller divergences, typically around 0.5%.

We observed that parsing ML trees built for HPV variants into branches containing established sublineage reference variants yielded lineages and sublineages assignments that largely – but not always - match ANI-based classification. Notably, HPV variants within the same lineage or sublineage have shared subtle evolutionarily conserved sequence features, such as lineage-specific and sublineage-specific SNPs detectable through multiple sequence alignment.

### Informative Positions in Frequency Profiles

Profile positions were classified as *trivial or non-trivial*. Trivial positions exhibit no nucleotide variation across all genomes of a given HPV type. We observed that the number of trivial positions decreases as the number of genomes included in the MSA increases. Non-trivial positions were further divided into high-informative or low-informative categories (Table 3). High-informative positions are characterized by different dominant nucleotides - that is, nucleotides with the highest frequency at a given position - across at least two sublineages. In contrast, low-informative positions share the same dominant nucleotide across all sublineages, but with substantially different nucleotide frequencies.

**Table 3.** Statistics on high- and low-informative positions for the nine HPV types.

| HPV type | # of nontrivial positions in profile | # of low informative positions | # of high informative positions |
| --- | --- | --- | --- |
| HPV 6 | 477 | 337 | 140 |
| HPV 11 | 261 | 112 | 149 |
| HPV 16 | 1,279 | 982 | 297 |
| HPV 18 | 647 | 381 | 266 |
| HPV 31 | 458 | 370 | 88 |
| HPV 33 | 243 | 89 | 154 |
| HPV 45 | 1,007 | 813 | 194 |
| HPV 52 | 727 | 482 | 245 |
| HPV 58 | 537 | 349 | 188 |

Locations of high-informative positions for HPV-58 at the lineage and sublineage levels are shown in Fig. 4A and Fig. 4B respectively. Locations of high-informative profile positions for lineages and sublineages of all nine HPV types are provided in Suppl. Fig. 3.

**Figure 4.**
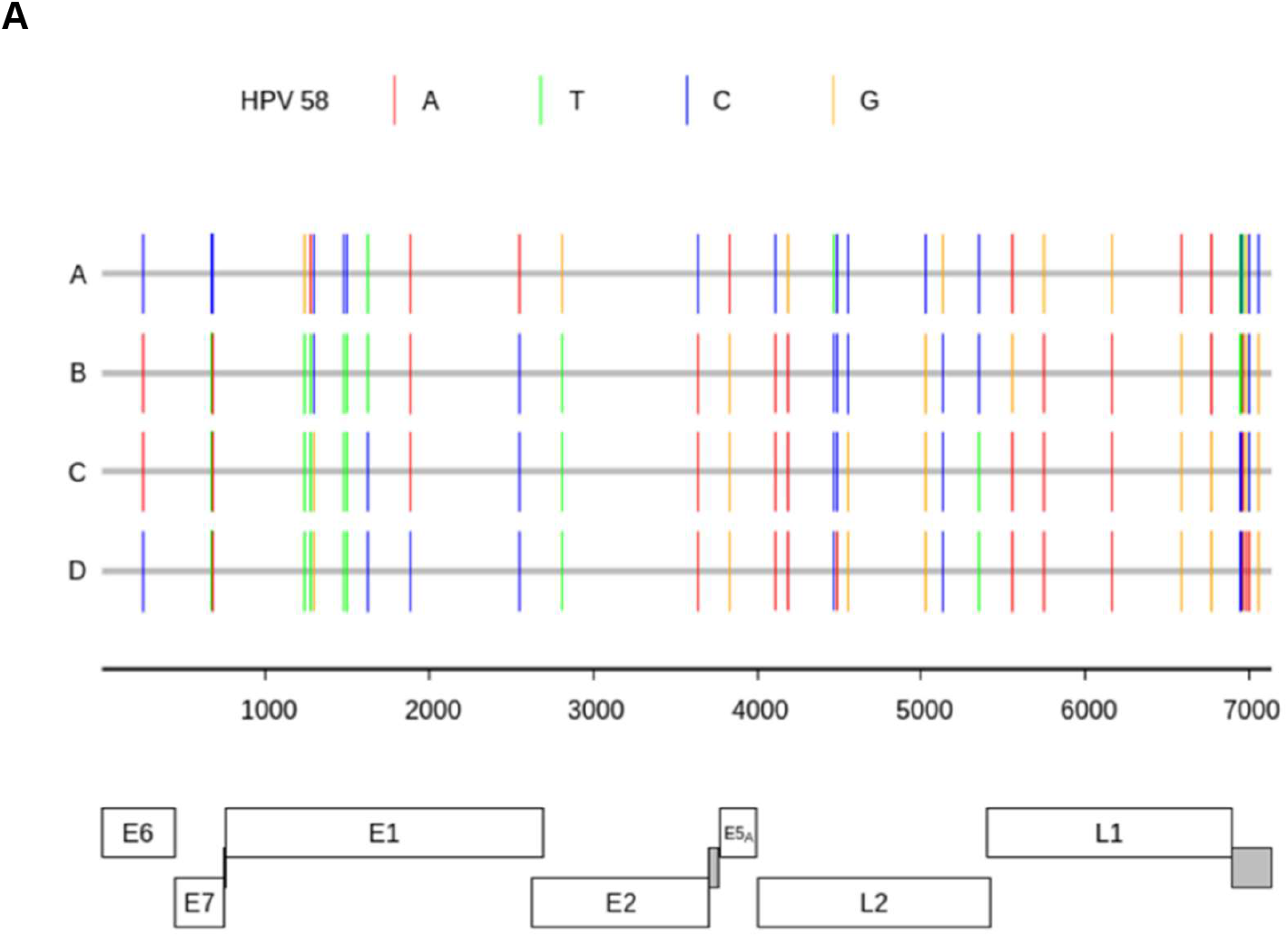
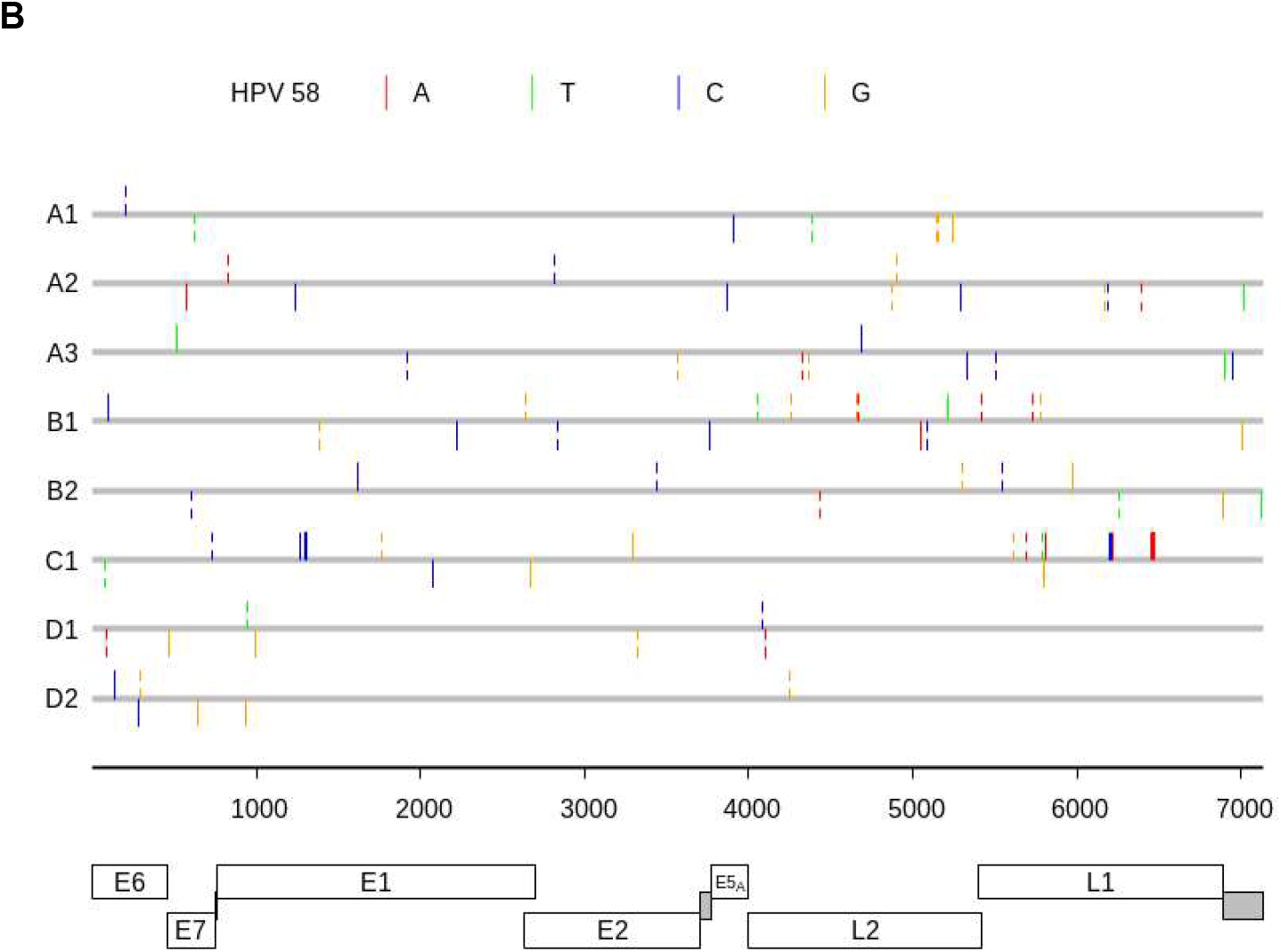
High-informative profile positions for the lineages (A) and for sublineages (B) of HPV-58. Broken ticks correspond to the third position of a codon. Whole ticks appear either in the first and second codon positions or in intergenic regions.

We observed that when the tested fragments overlapped with high-informative positions classification error rates decrease substantially. However, low-informative positions also contributed to making the profile-based predictions more precise.

### Tests of the Algorithm for Variant Calling on Partial HPV Genome Sequences

Tests of the new algorithm predicting the taxonomic affiliation of a query fragment at the lineage and sublineage level were performed using fragments of 1,000 nt and 2,000 nt taken from genomes with known taxonomic affiliation. As could be expected, increasing fragment length reduced the classification error rate (Table 4). For all HPV types except HPV-16 and HPV-31, the error rate dropped below 1% for the 2,000 nt fragments. We also tested classification accuracy using even longer fragments. Error rates decreased steadily as fragment length increased (data not shown).

**Table 4.** Error rates (%) observed in predicting the taxonomic origin of 4,000 randomly selected fragments of each HPV type. The error rates are listed for taxonomic classification by the profile posterior probability score (panel A) and for classification by the highest BLASTN score (panel B).

| HPV type | 1,000 nt |  | 2,000 nt |  |
| --- | --- | --- | --- | --- |
|  | lineage | sublineage | lineage | sublineage |
| 6 | 1.9 | 6.6 | 0.5 | 1.6 |
| 11 | 0.0 | 6.4 | 0.0 | 0.4 |
| 16 | 0.1 | 18.8 | 0.2 | 9.1 |
| 18 | 0.0 | 3.6 | 0.0 | 0.2 |
| 31 | 5.4 | 17.1 | 0.0 | 3.1 |
| 33 | 0.1 | 0.3 | 0.0 | 0.0 |
| 45 | 2.1 | 5.0 | 0.1 | 1.3 |
| 52 | 4.2 | 9.7 | 0.7 | 1.4 |
| 58 | 1.6 | 5.7 | 0.0 | 0.1 |

**B**
| HPV type | 1,000 nt |  | 2,000 nt |  |
| --- | --- | --- | --- | --- |
|  | lineage | sublineage | lineage | sublineage |
| 6 | 0.9 | 14.9 | 0.1 | 12.4 |
| 11 | 0.0 | 27.4 | 0.0 | 16.9 |
| 16 | 0.1 | 30.5 | 0.1 | 18.5 |
| 18 | 0.1 | 25.3 | 0.0 | 12.0 |
| 31 | 4.1 | 33.3 | 1.6 | 27.4 |
| 33 | 0.8 | 4.4 | 0.7 | 2.4 |
| 45 | 5.2 | 14.8 | 1.2 | 6.5 |
| 52 | 1.9 | 13.4 | 0.6 | 3.7 |
| 58 | 1.4 | 4.8 | 0.1 | 1.8 |

For comparison, we also predicted the taxonomic affiliation of query fragments using BLASTN alignments to sublineage reference genomes. The sublineage corresponding to the highest BLASTN alignment score was assigned to the fragment.

Overall, the results demonstrate a clear advantage of the profile-based method, which yields lower error rates - especially when distinguishing between sublineages belonging to the same lineage.

To confirm that the 4,000 fragment test sets were sufficient for evaluating performance, we repeated the analysis using 50,000 fragments sets. Comparison of the error rates indicates that the smaller test sets provide estimates close to the rates determined for the large set of fragments (Suppl. Table 6).

We also carried out a more detailed analysis of the error rates for groups of fragments originating from the same genomic region. The error rate varied across genomic locations, and this variation correlated with the number of informative positions contained within a query fragment (data not shown).

The relatively high overall error rates for HPV-16 and HPV-31 can be explained by the structure of the ANI divergence distributions. In HPV-16 all but one ANI divergence between four sublineages of lineage C and four sublineages of lineage D are less than 1.0 – same for sublineages of lineages A and B for HPV-31 – which comes in contrast with other HPV types (and other pairs of lineages in HPV-16 and HPV-31) where the intra-lineage distances between sublineages are larger than 1.0 and sometimes rise above 2.0 (bold font numbers in Suppl. Table 5). Classification of fragments that are very similar to sequences from other sublineages is inherently more error-prone than classification of fragments that are more clearly separated in sequence space.

### Concluding Remarks

Our aim was to develop an algorithm that assigns taxonomic labels – lineages and sublineages - to DNA sequences obtained from clinical or epidemiological HPV studies. A conventional approach to such classification is to incorporate a new (often partial) sequence into a multiple sequence alignment, reconstruct a phylogenetic tree, and infer the placement of the query sequence (Mirabello et al. 2016). However, tree-based classification is computationally expensive, as the entire tree must be rebuilt each time a new sequence is added.

A faster alternative is to align the query fragment pairwise to reference sublineage genomes and assign it to the sublineage yielding the best alignment score. However, we have shown that the accuracy of such method is not high enough (Table 4B).

The proposed profile-based method requires a one-time training step, with a runtime comparable to that of a single tree-based classification (Burk, Harari, and Chen 2013). A substantial reduction in training time is achieved by replacing CLUSTALO (Sievers and Higgins 2021) with the fast-MSA algorithm developed specifically for closely related HPV genomes.

Once training is complete, classification of individual query sequence is extremely fast - comparable to pairwise alignment methods – while offering improved sensitivity by leveraging information aggregated from multiple aligned genomes rather than a single representative. The profile-based method requires less than one second per sequence, enabling the classification of approximately 3,600 sequences in under an hour. In contrast, CLUSTALO, followed by RAxML requires more than five hours to construct MSAs and ML trees for 1,000 sequences.

Finally, we emphasize the central role of HPV reference genomes in this framework. Reference genomes of HPV sublineages, curated and proposed by groups of experts in HPV evolution, are critical both for sublineage clustering during tree parsing and for parameterizing the sublineage-specific probabilistic profiles used in our method.

## Software availability

The HPV variant calling software, HPVarcall, is available on GitHub at https://github.com/gatech-genemark/HPVarcall). All scripts and datasets used to generate the figures and tables in this paper are available at https://github.com/gatech-genemark/HPVarcall-exp.

## Acknowledgement

We are grateful to Elizabeth Unger and Mangalathu Rajeevan for valuable comments and suggestions. This work was supported in part by a grant AWD-006572 from the US Centers for Disease Control and Prevention.

## Notes

### Competing Interest Statement

The authors have declared no competing interest.

