## Supplemental_Figures for "HPVarcall: Calling lineages and sublineages for partial DNA sequences of human papillomavirus"

Supplemental Figure 1

ML trees for genomes of the nine HPV types

(Each type of HPV has its own specific panel.)

HPV 6

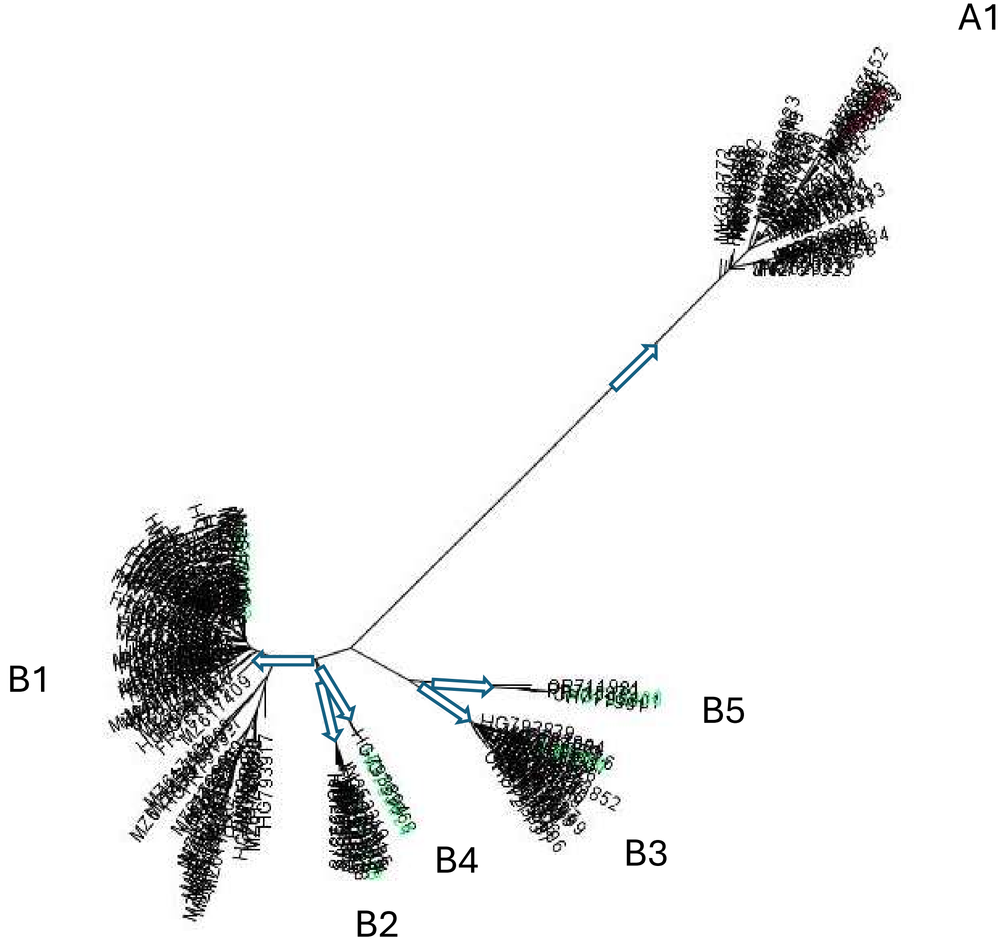

HPV 11

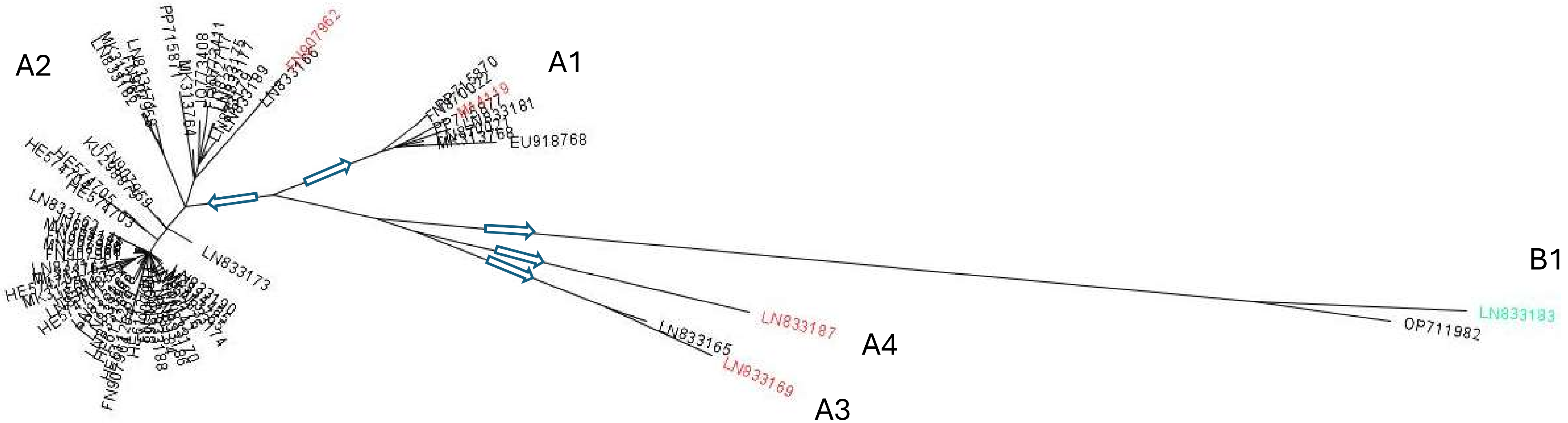

A2
A1
B1
A4
A3
FN907962
EU918768
LN833181
LN833173
LN833187
LN833165
LN833169
LN833183
OP711982

HPV 16

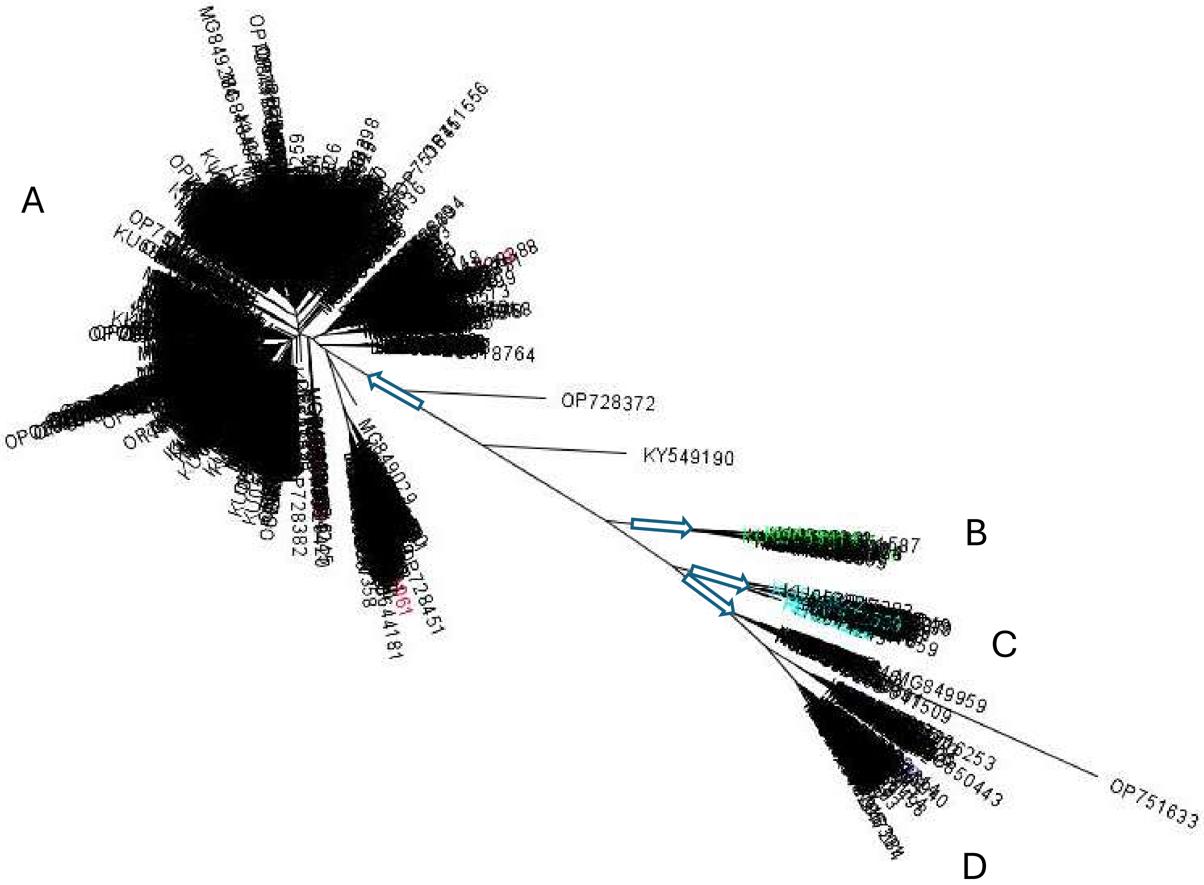

HPV 18

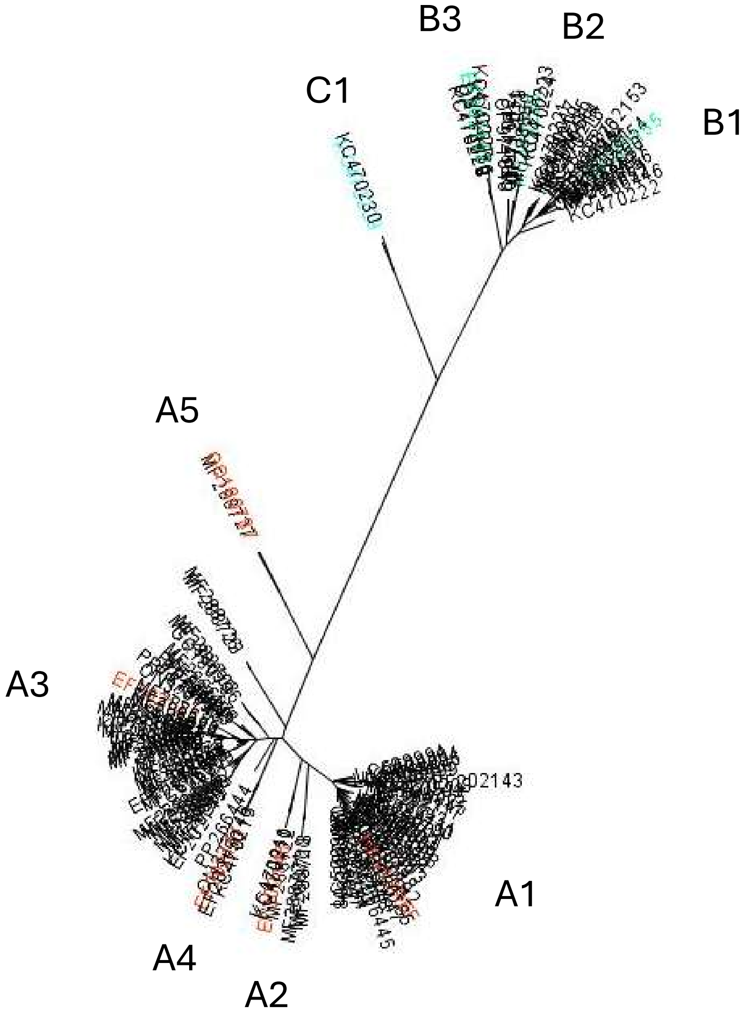

HPV 31

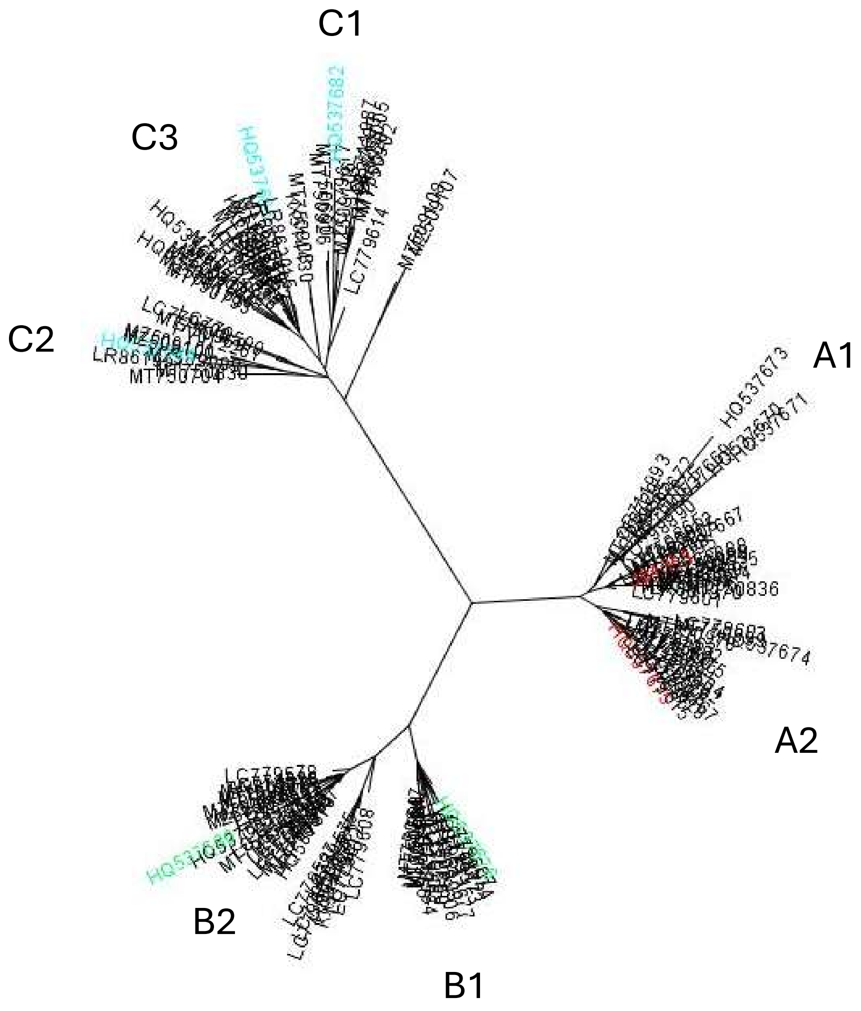
C1
C3
C2
A1
A2
B2
B1
HQ537673
HQ537671
HQ537674
LC779614
LC779678
HQ537682

HPV 33

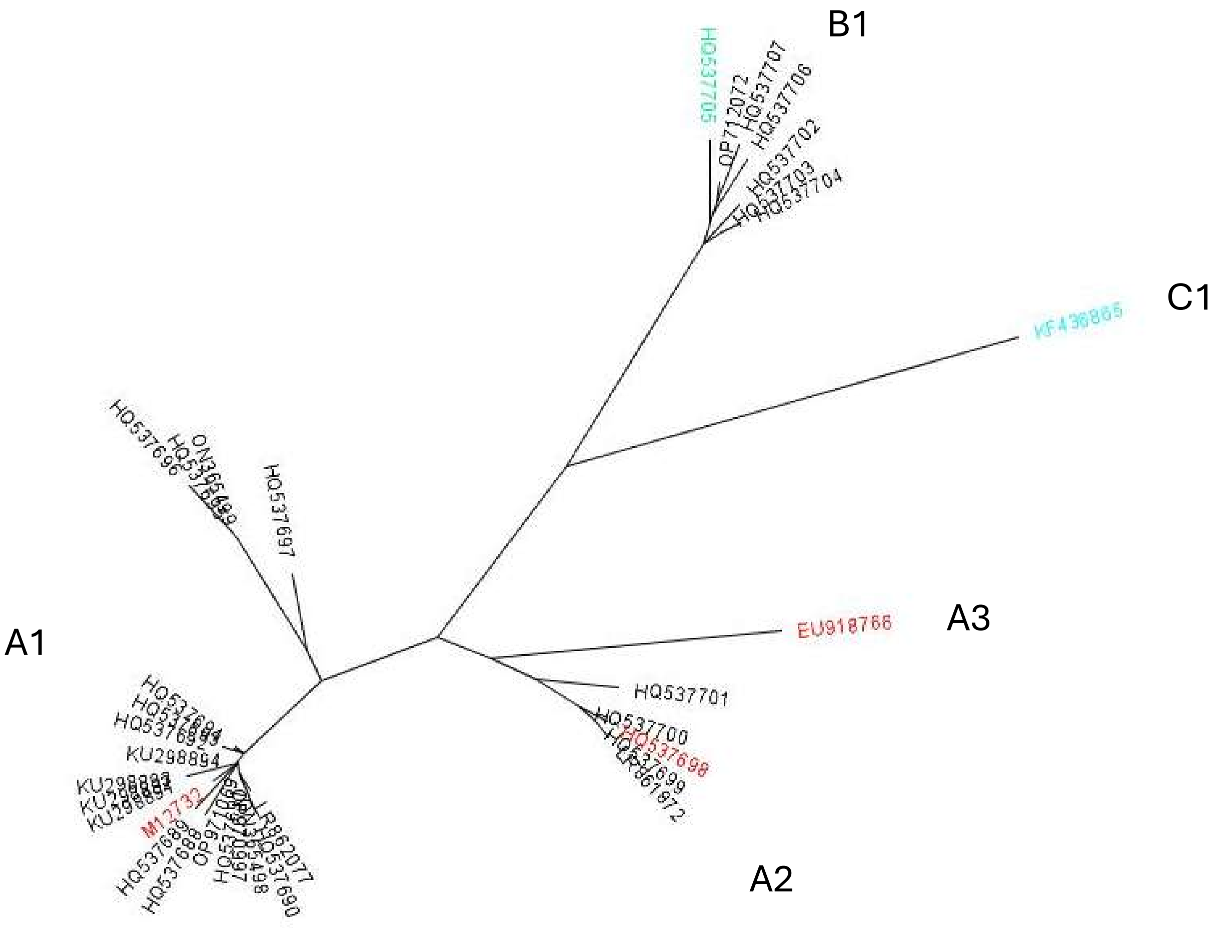

B1
HQ537705
HQ537707
OP712072
HQ537706
HQ537702
HQ537704
C1
KF436865
A3
EU918766
HQ537701
HQ537700
HQ537698
A2
A1
HQ537697
HQ537696
KU298894
M12732
LR862077
HQ537690
HQ537688
HQ537689

HPV 45

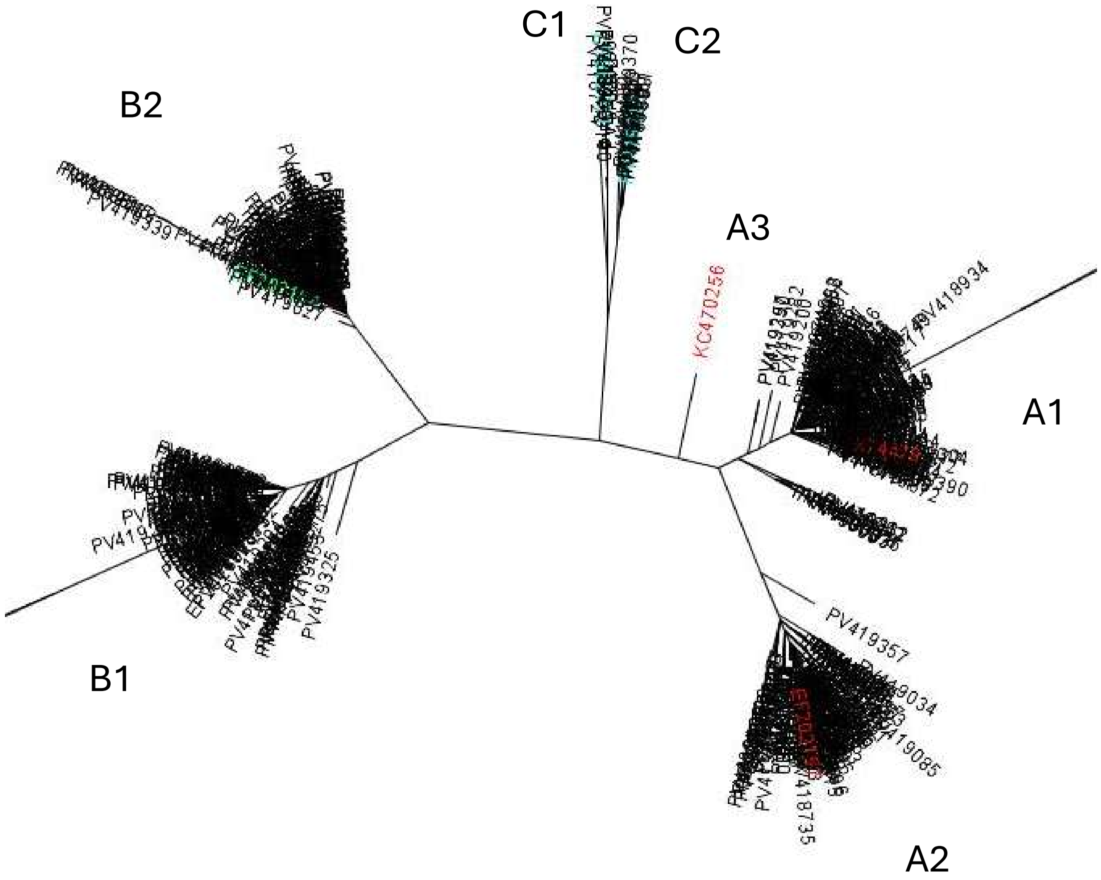

HPV 52

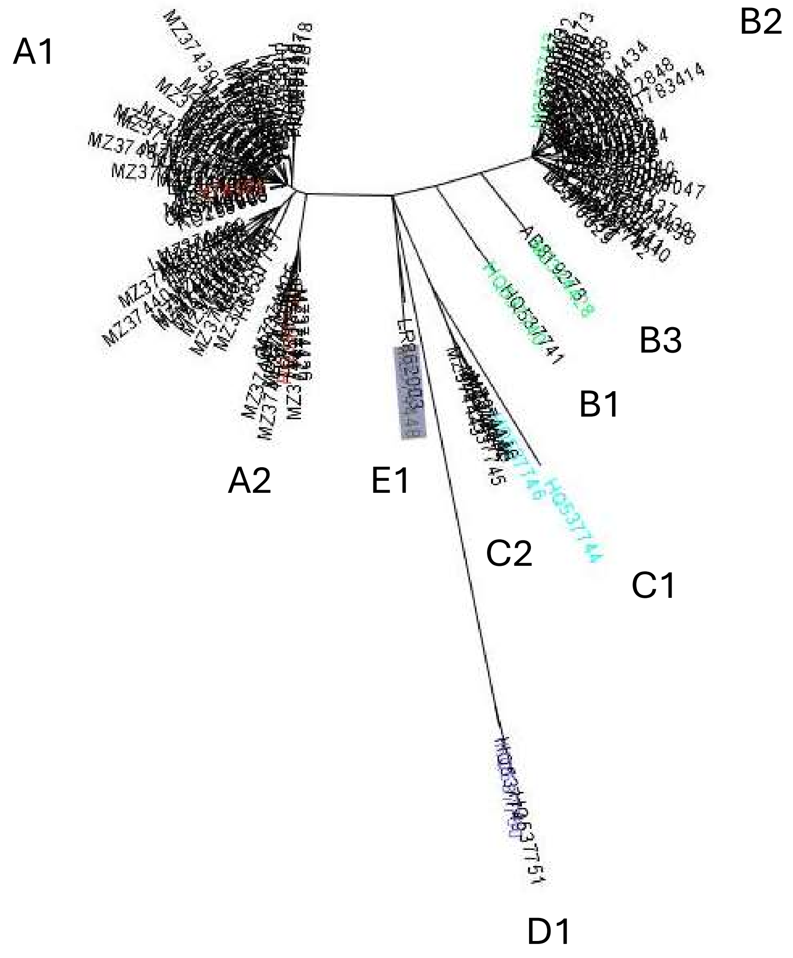

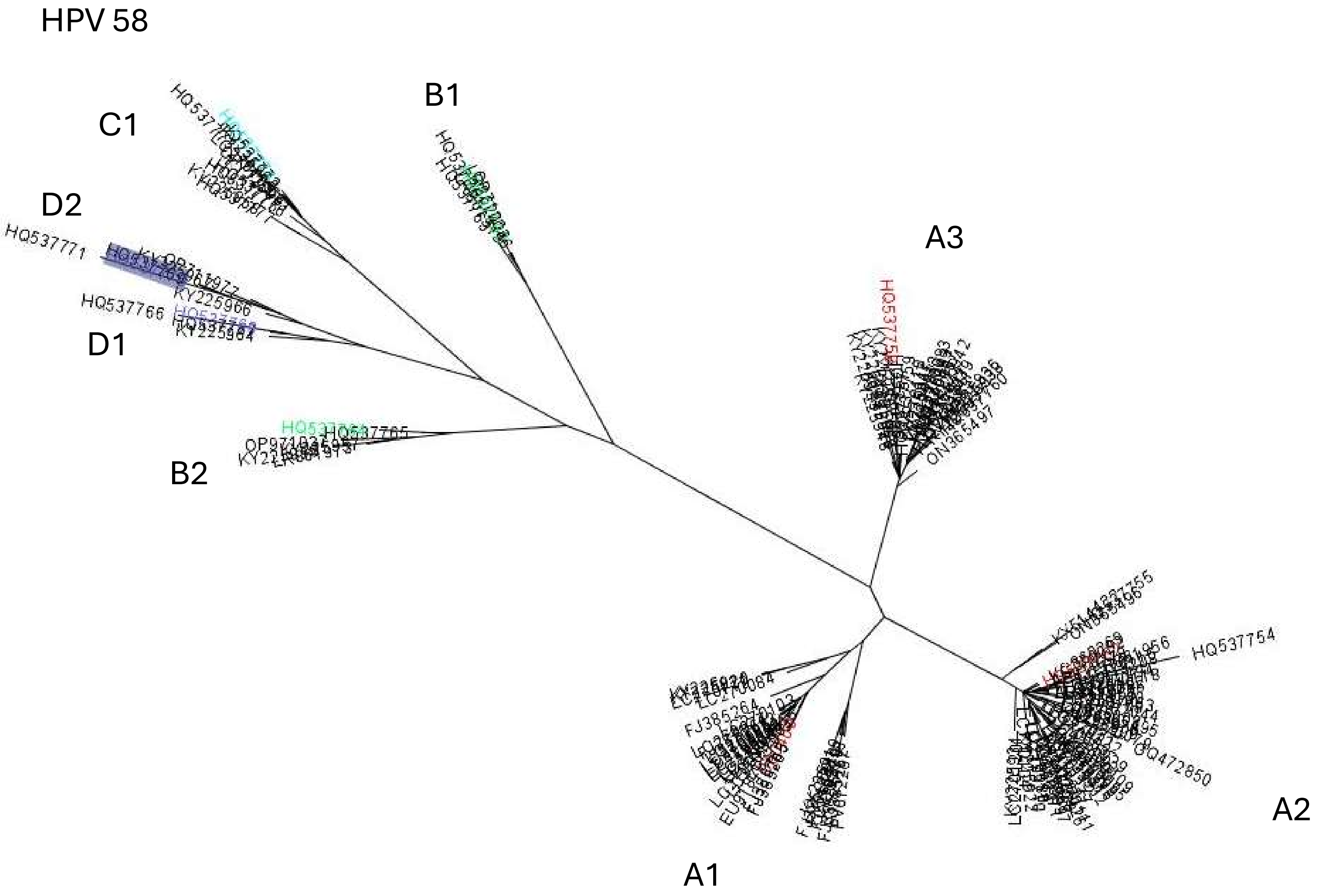
HPV 58
C1
B1
D2
HQ537771
KY225966
HQ537766
KY225964
D1
B2
A3
ON365497
HQ537754
GQ472850
FJ385264
A2
A1

Supplemental Figure 2

A -Genome-wide distribution of **lineage**-specific high-informative SNP positions for each of the nine HPV types.
(in the next nine panels)

B -Genome-wide distribution of **sublineage**-specific high-informative SNP positions for each of the nine HPV types.
(in the subsequent nine panels)

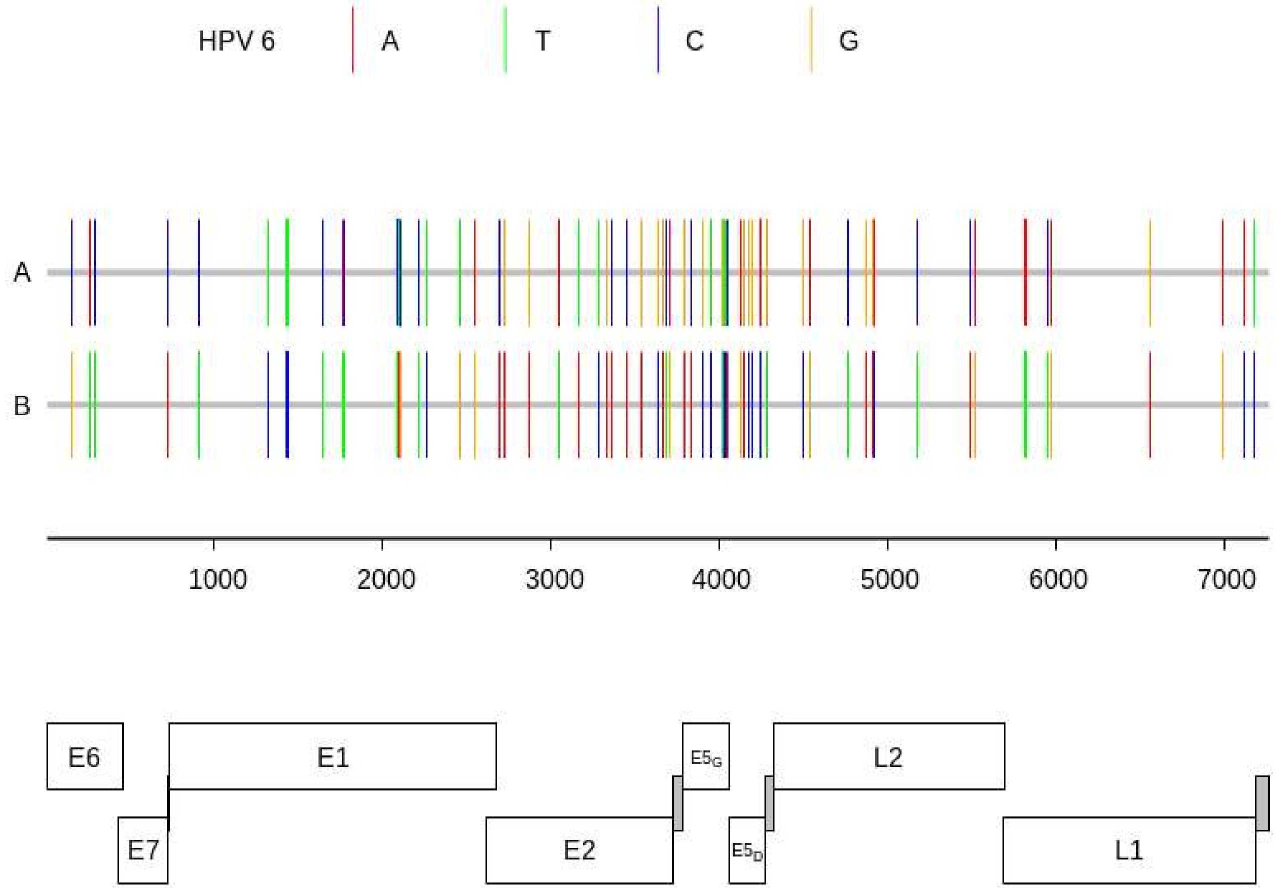

HPV 6
A
T
C
G
A
B
1000
2000
3000
4000
5000
6000
7000
E6
E7
E1
E2
E5G
E5D
L2
L1

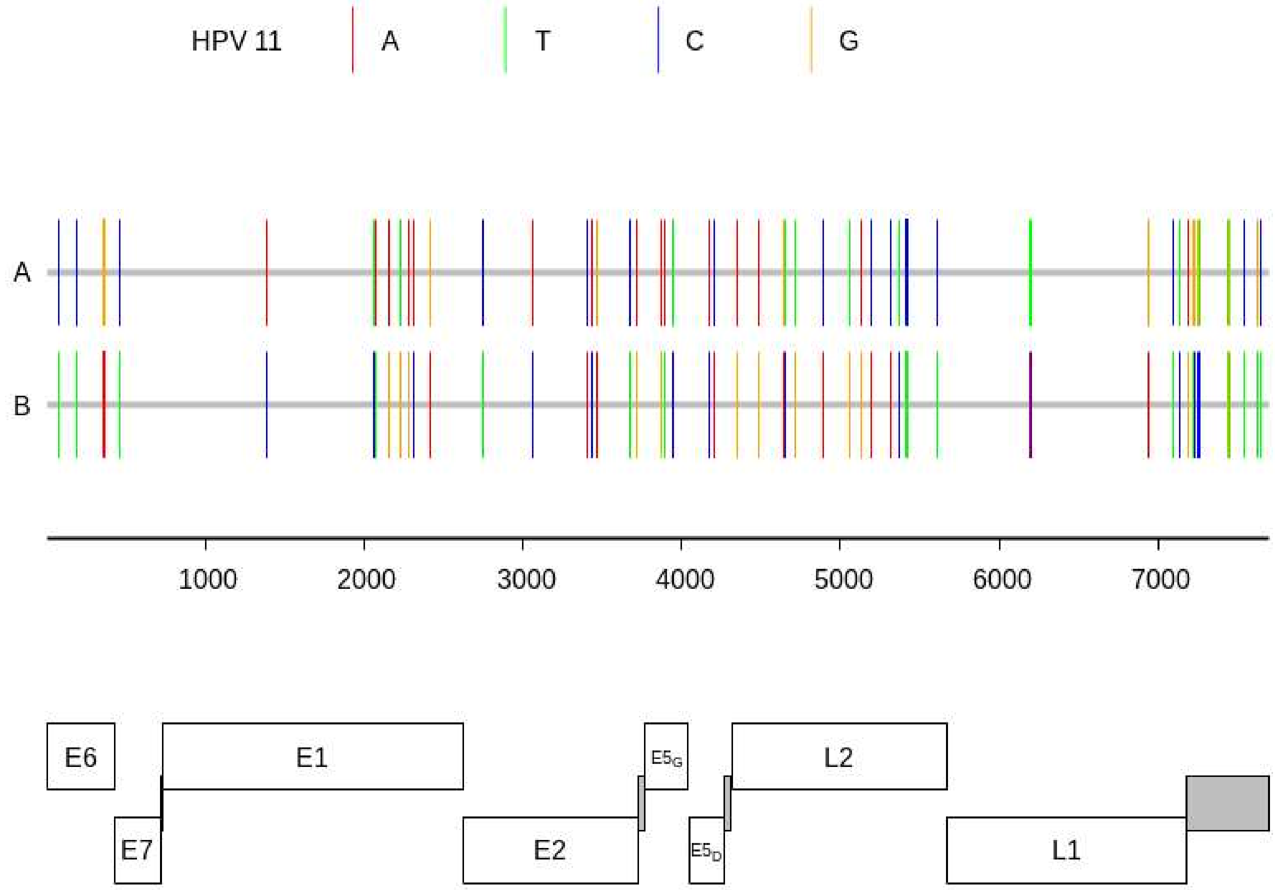

HPV 11
A
T
C
G
A
B
1000
2000
3000
4000
5000
6000
7000
E6
E1
E5G
L2
E7
E2
E5D
L1

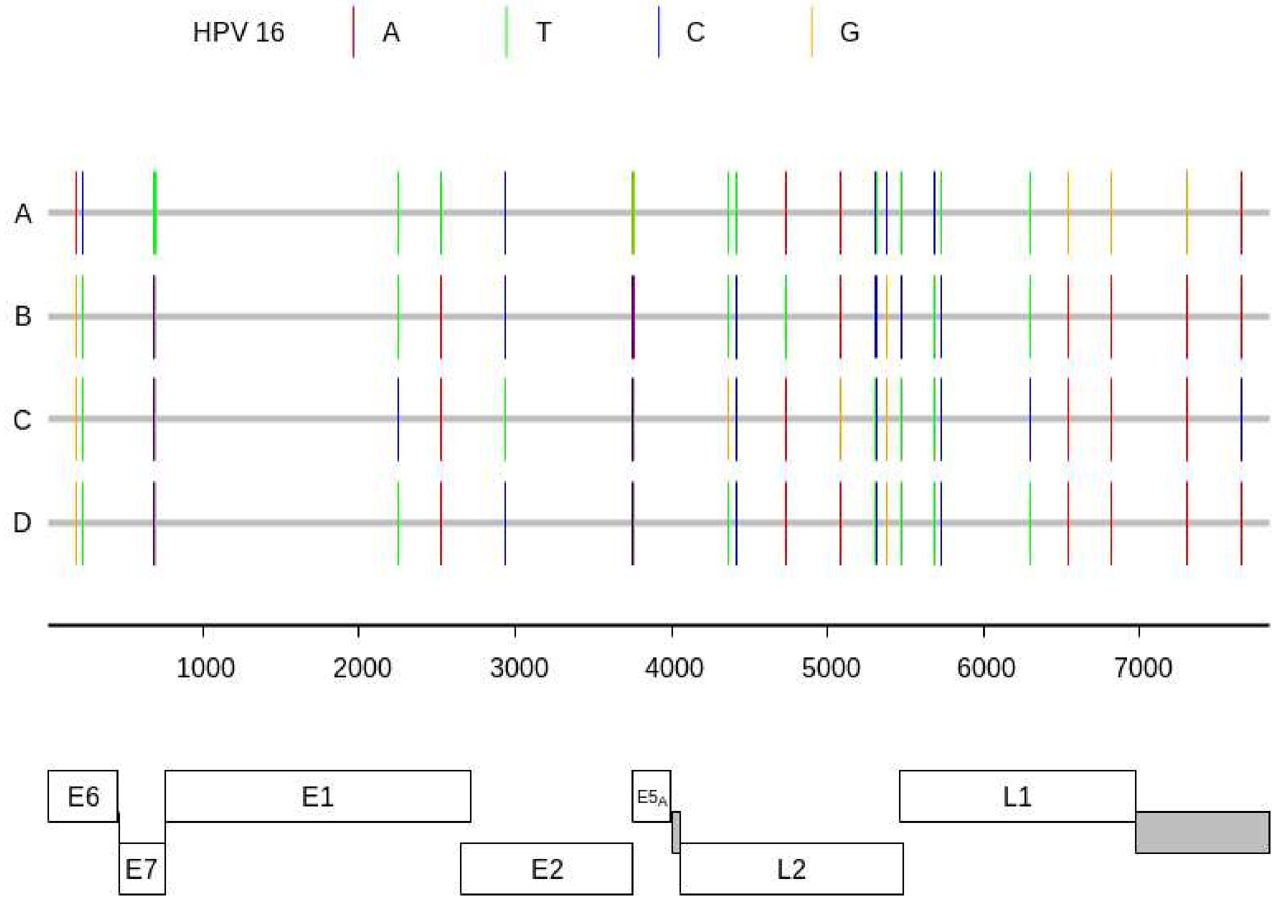

HPV 16
A
T
C
G
A
B
C
D
1000
2000
3000
4000
5000
6000
7000
E6
E7
E1
E2
E5A
L2
L1

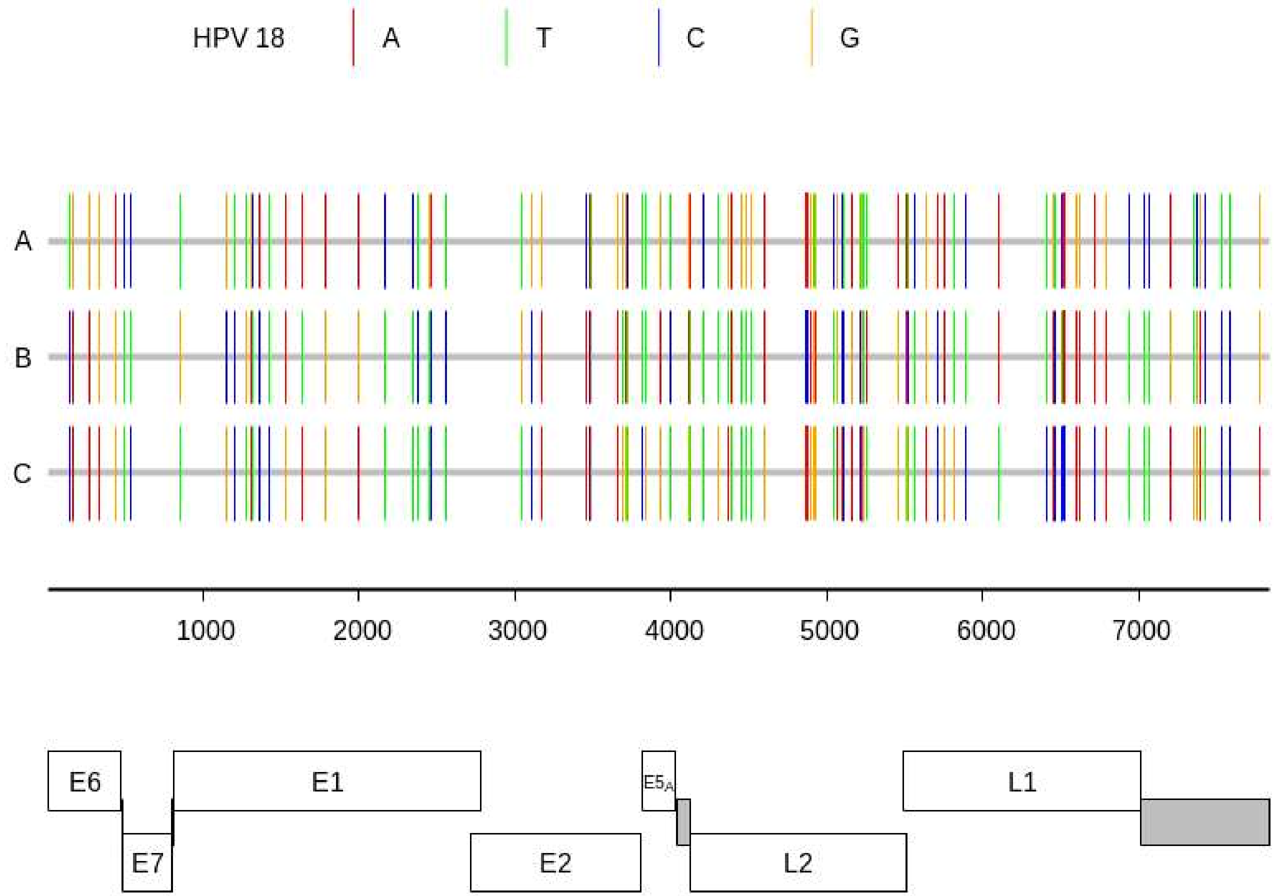

HPV 18
A
T
C
G
A
B
C
1000
2000
3000
4000
5000
6000
7000
E6
E7
E1
E2
$E5_A$
L2
L1

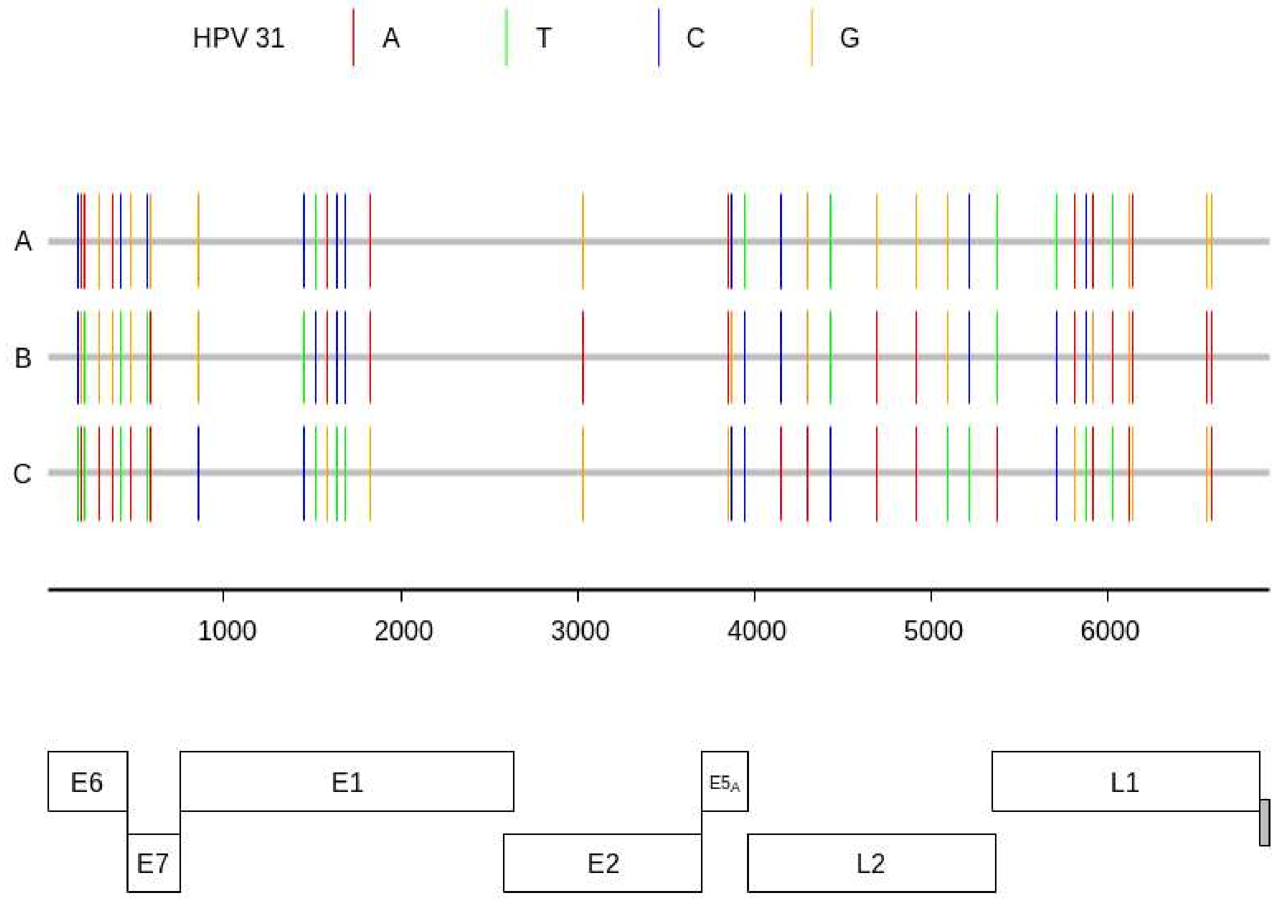

HPV 31
A
T
C
G
A
B
C
1000
2000
3000
4000
5000
6000
E6
E7
E1
E2
E5A
L2
L1

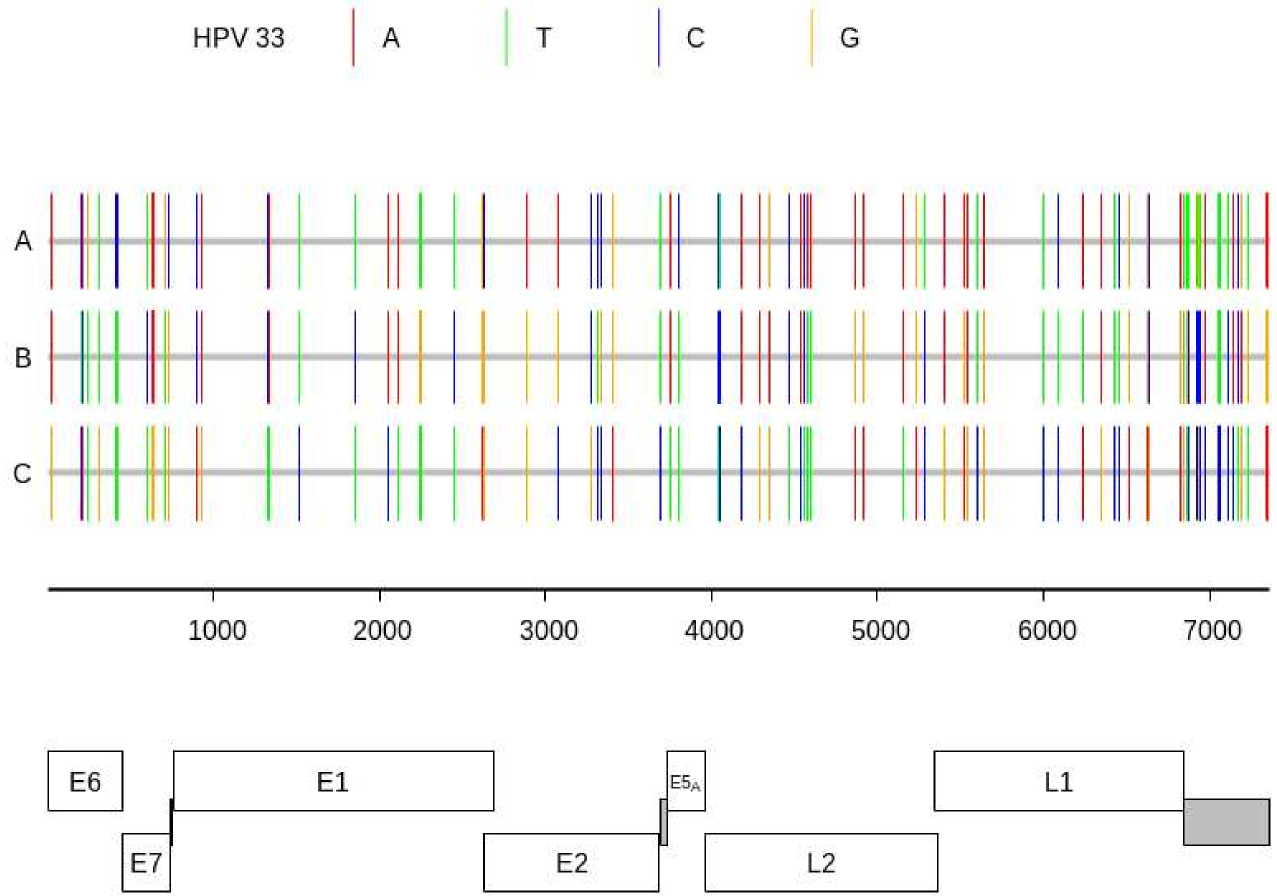

HPV 33
A
T
C
G
A
B
C
1000
2000
3000
4000
5000
6000
7000
E6
E7
E1
E2
E5A
L2
L1

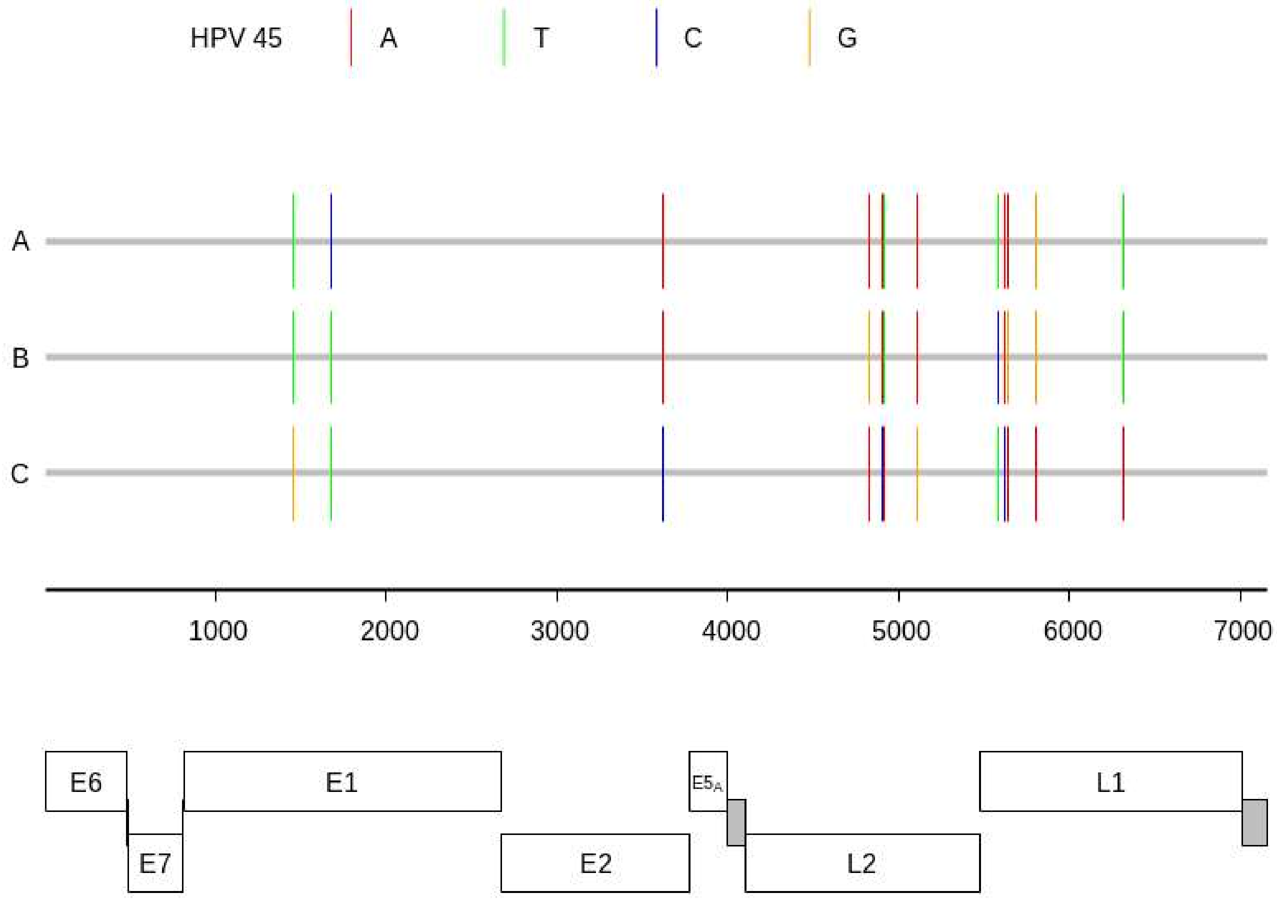
HPV 45
A
T
C
G
A
B
C
1000
2000
3000
4000
5000
6000
7000
E6
E7
E1
E2
E5A
L2
L1

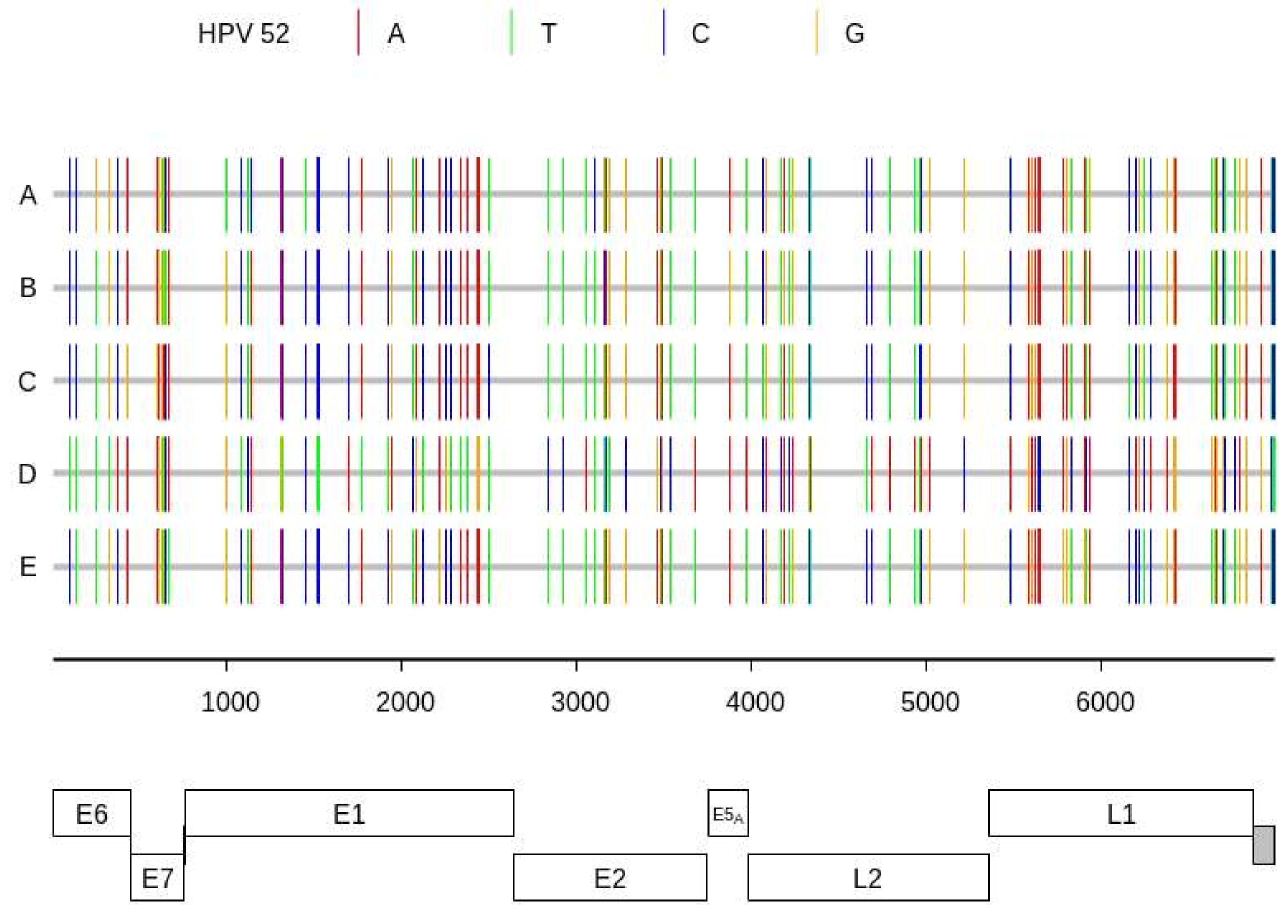

HPV 52
A
T
C
G
A
B
C
D
E
1000
2000
3000
4000
5000
6000
E6
E7
E1
E2
E5A
L2
L1

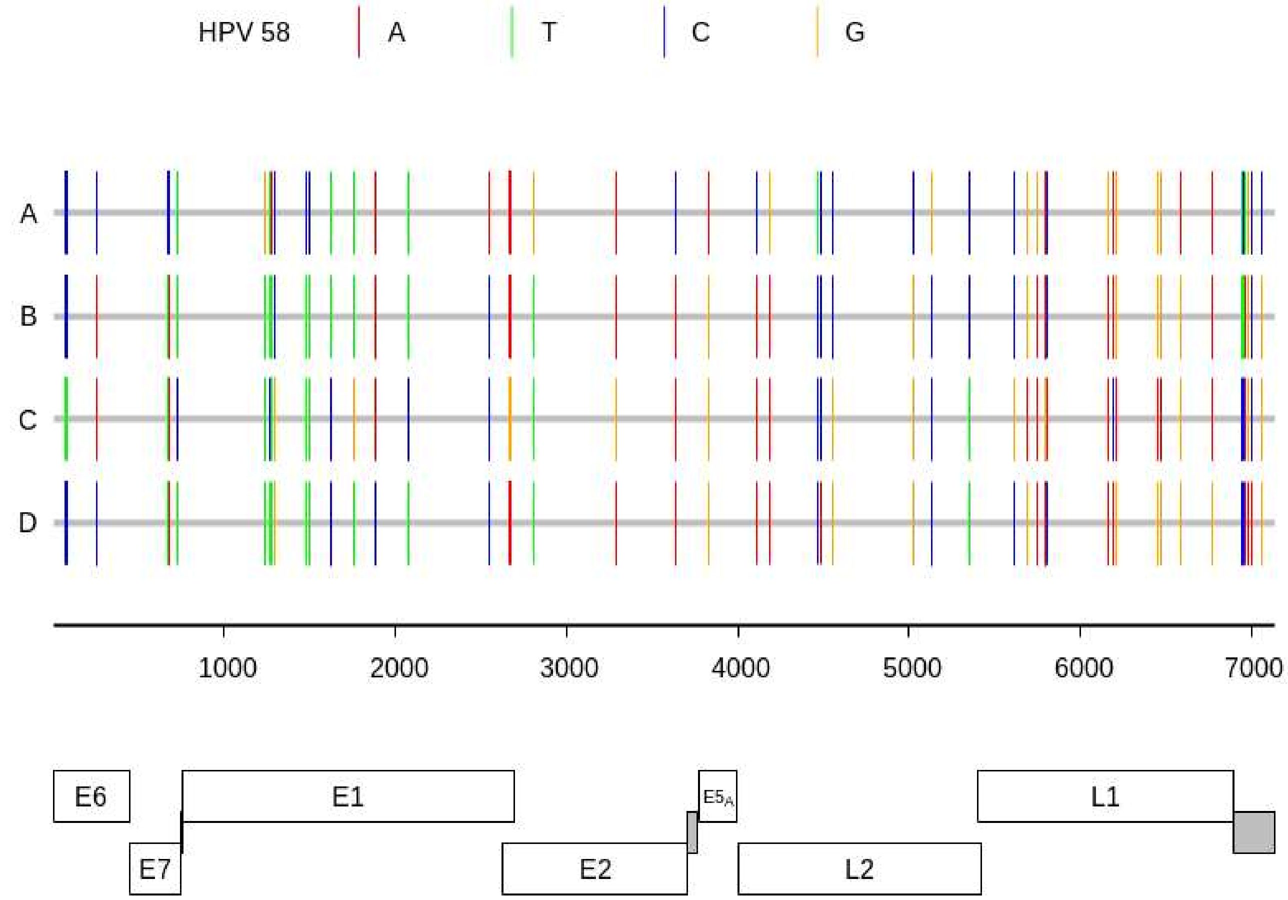

HPV 58
A
T
C
G
A
B
C
D
1000
2000
3000
4000
5000
6000
7000
E6
E7
E1
E2
E5A
L2
L1

B - Genome-wide distribution of **sublineage**-specific high-informative SNP positions for each of the nine HPV types. (in the next nine panels)

Broken ticks correspond to the third position of a codon.
Whole ticks appear either in the first and second codon positions or in intergenic regions

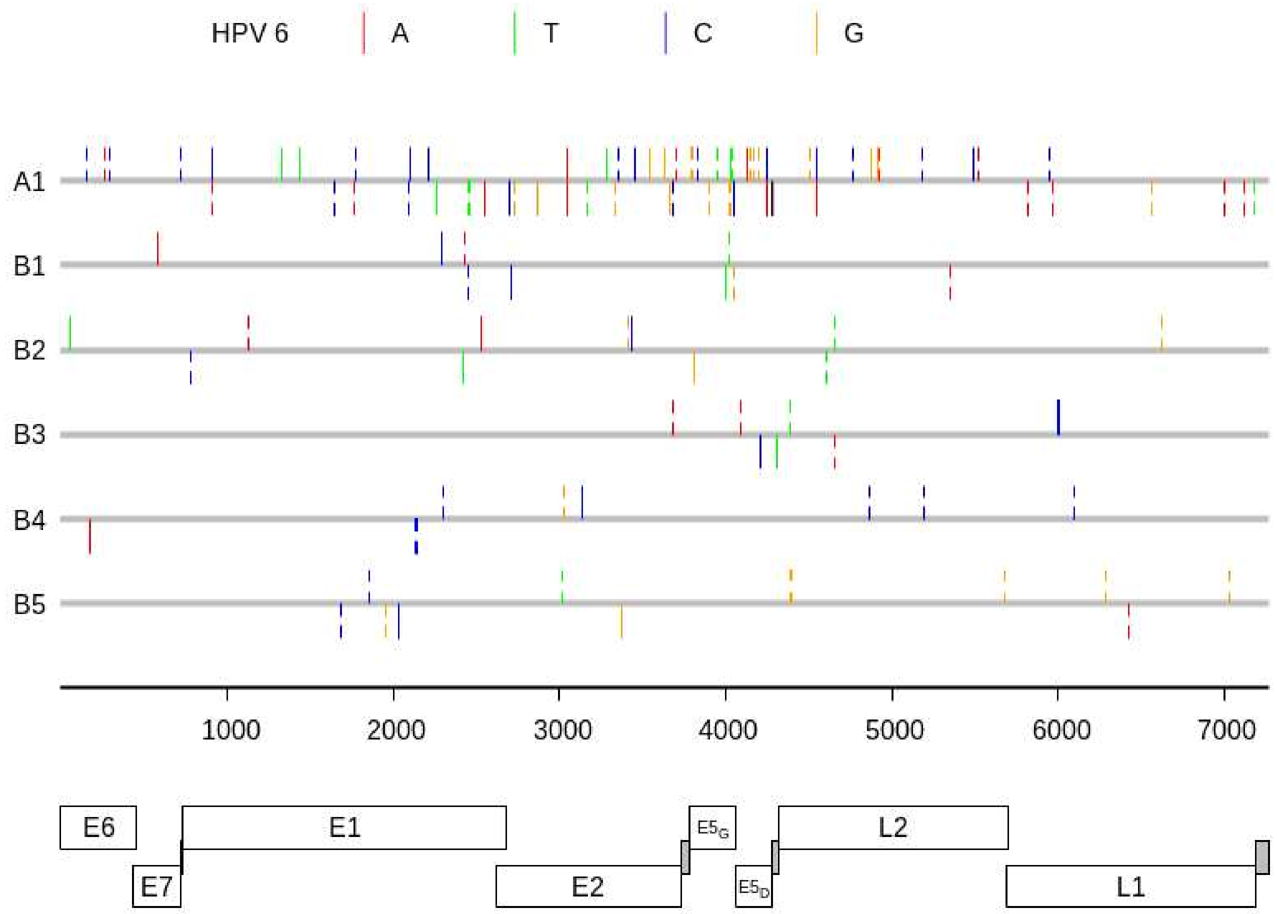

HPV 6
A
T
C
G
A1
B1
B2
B3
B4
B5
1000
2000
3000
4000
5000
6000
7000
E6
E7
E1
E2
E5G
E5D
L2
L1

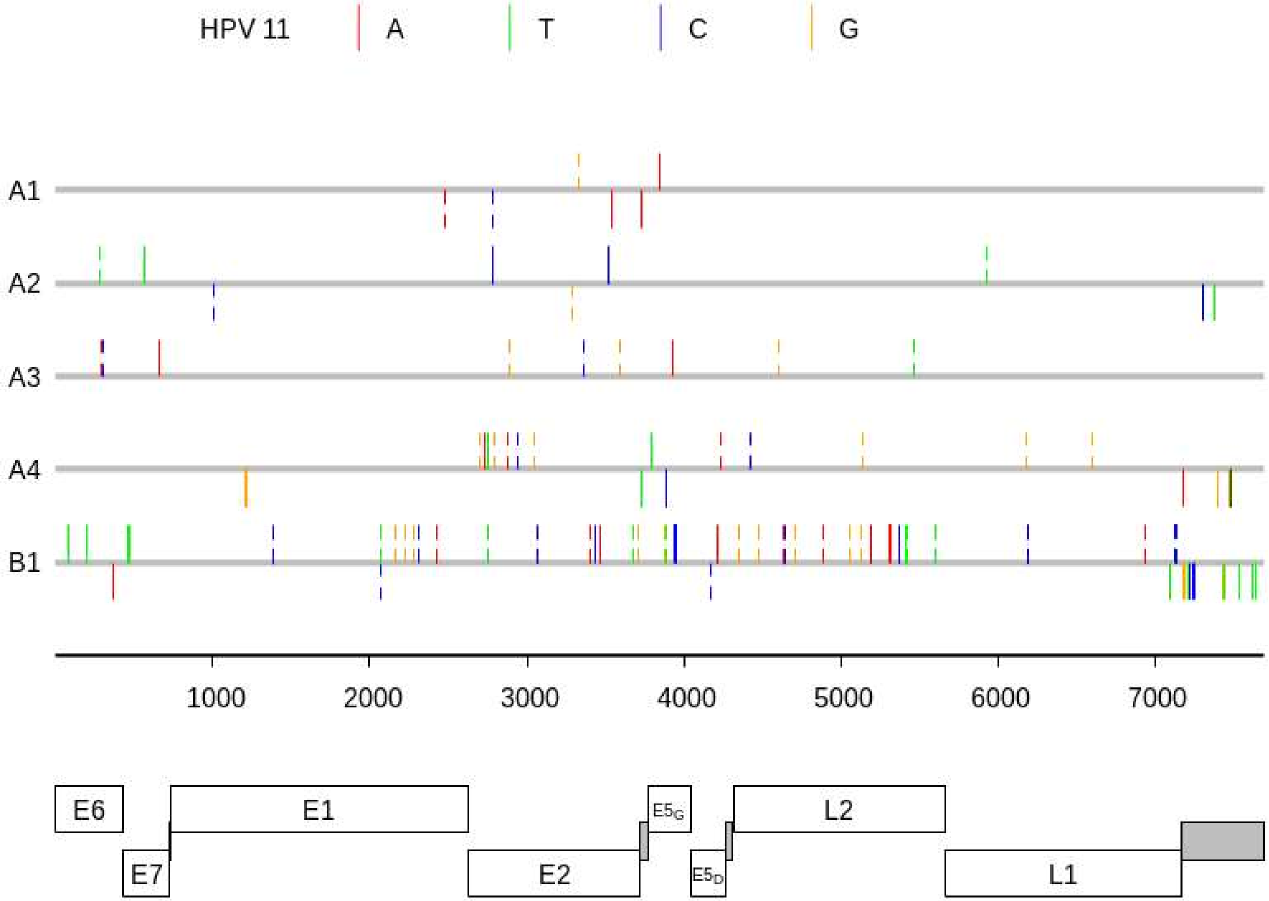

HPV 11
A
T
C
G
A1
A2
A3
A4
B1
1000
2000
3000
4000
5000
6000
7000
E6
E1
E5G
L2
E7
E2
E5D
L1

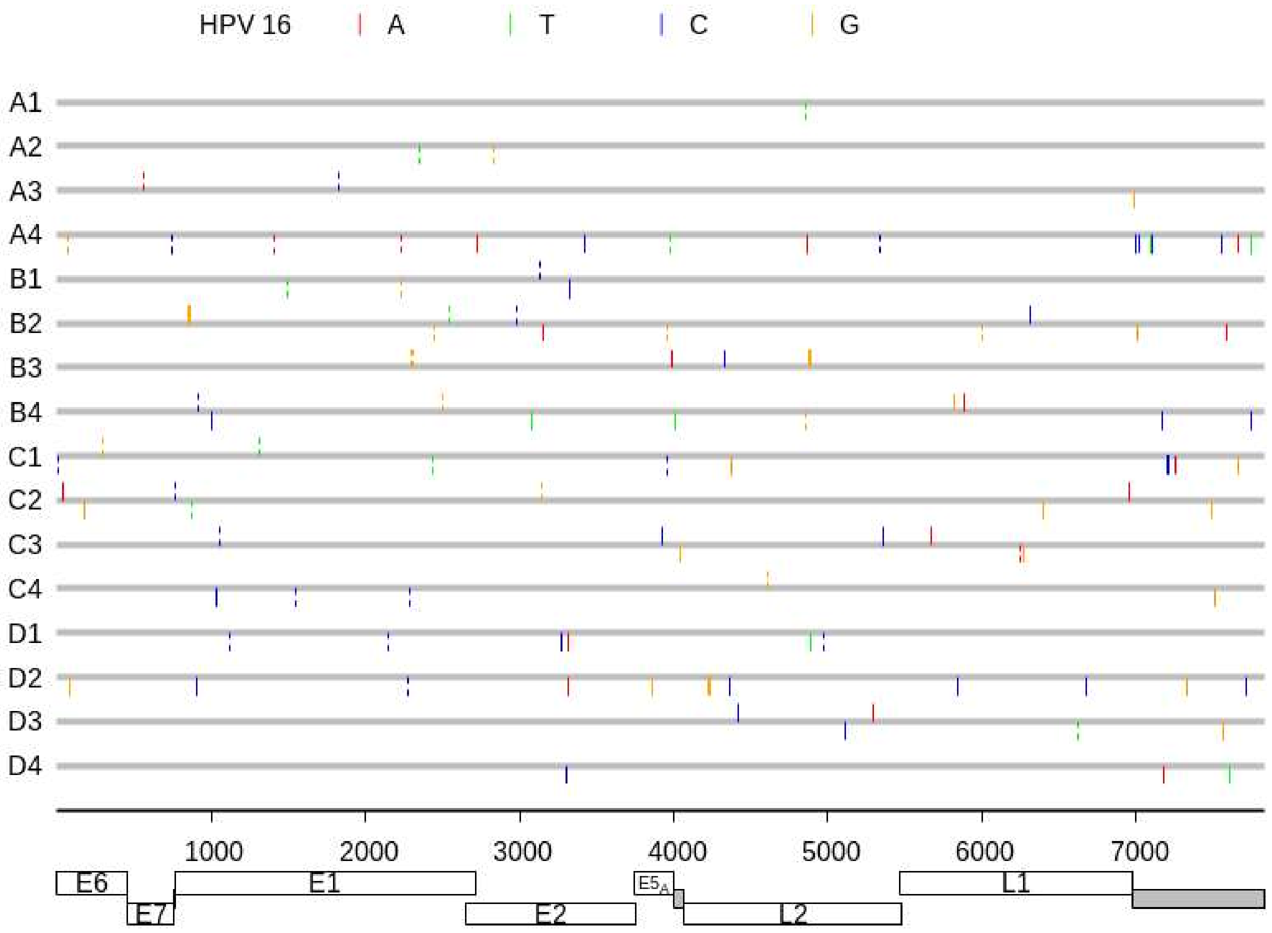

HPV 16
A
T
C
G
A1
A2
A3
A4
B1
B2
B3
B4
C1
C2
C3
C4
D1
D2
D3
D4
1000
2000
3000
4000
5000
6000
7000
E6
E7
E1
E2
E5A
L2
L1

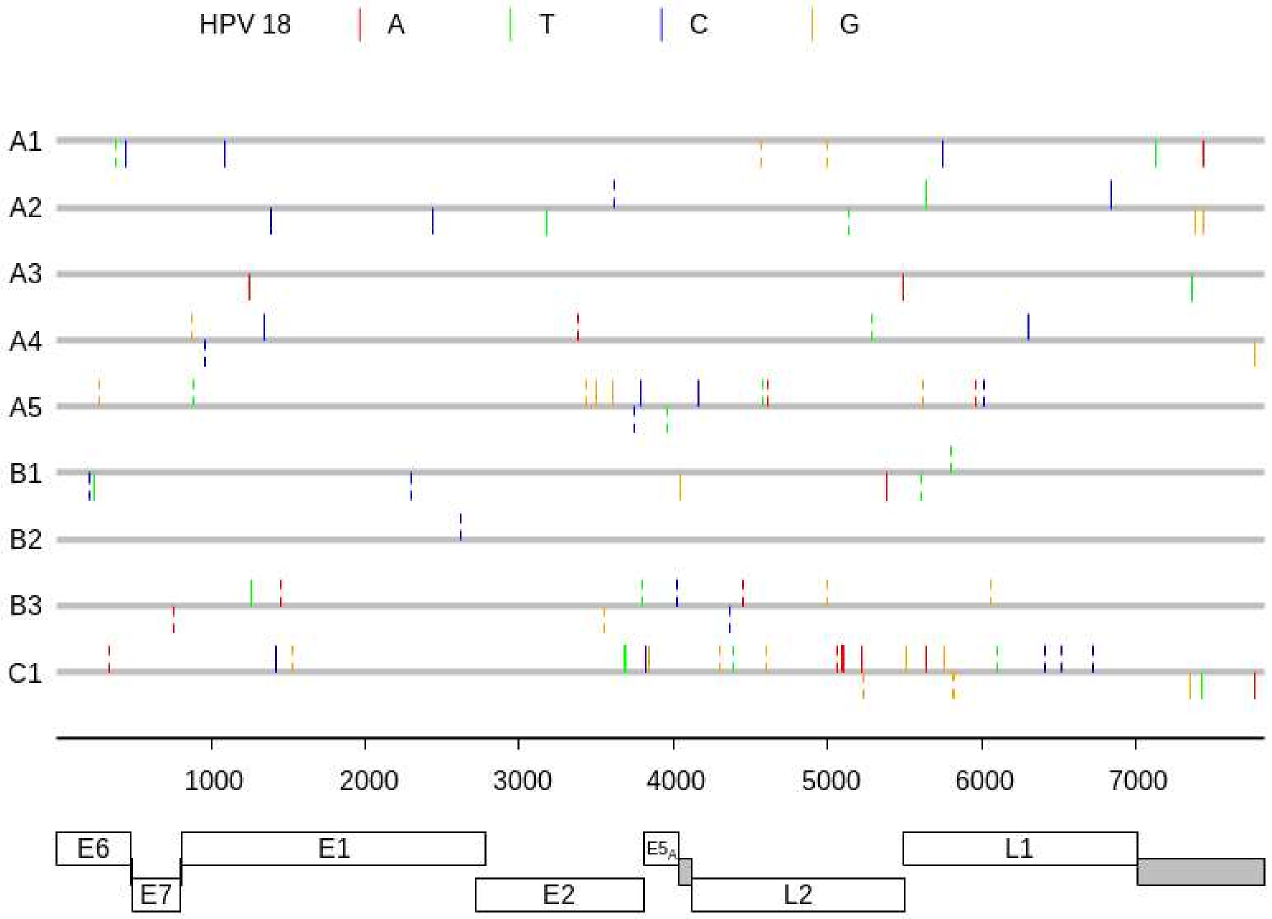

HPV 18
A
T
C
G
A1
A2
A3
A4
A5
B1
B2
B3
C1
1000
2000
3000
4000
5000
6000
7000
E6
E7
E1
E2
E5A
L2
L1

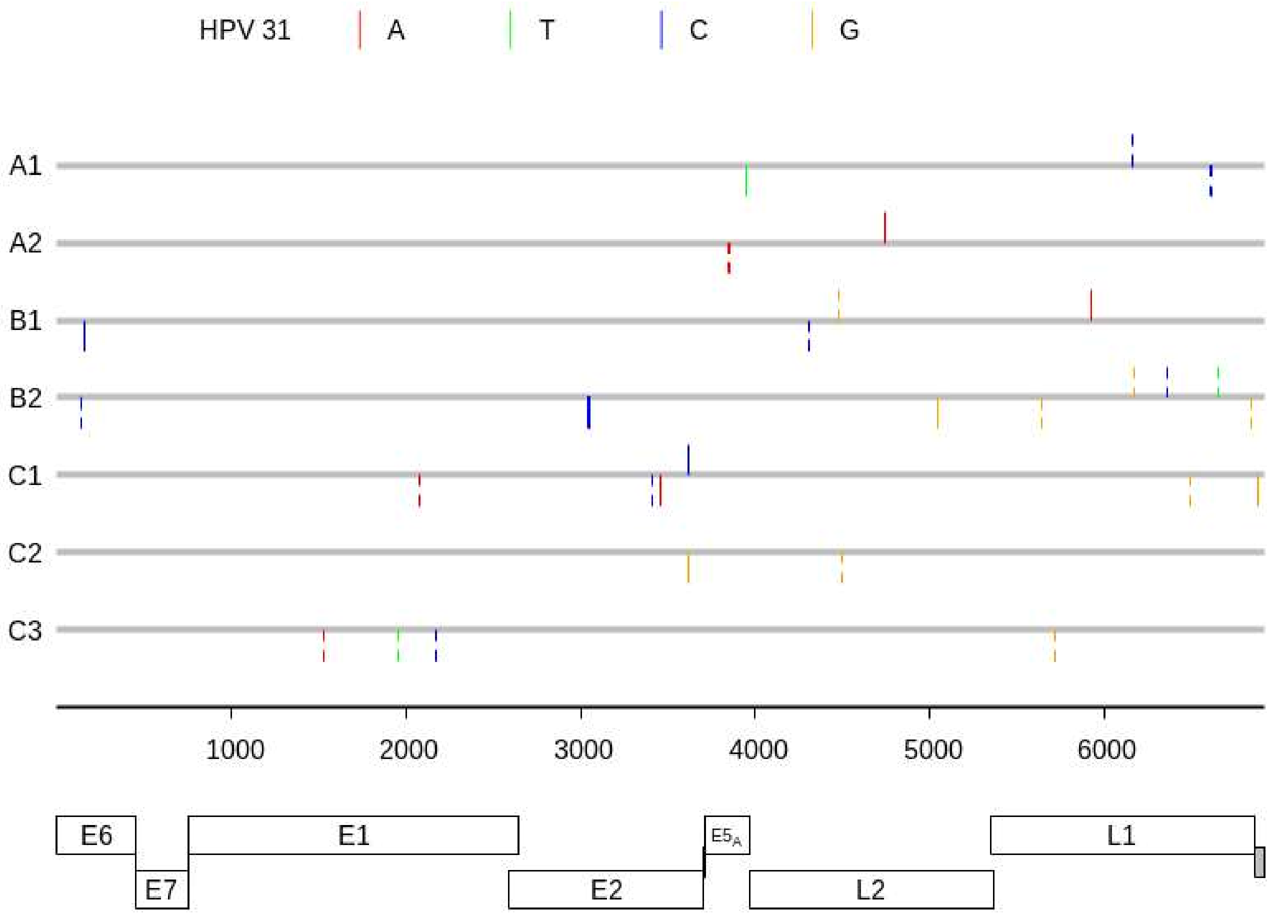

HPV 31
A
T
C
G
A1
A2
B1
B2
C1
C2
C3
1000
2000
3000
4000
5000
6000
E6
E7
E1
E2
E5A
L2
L1

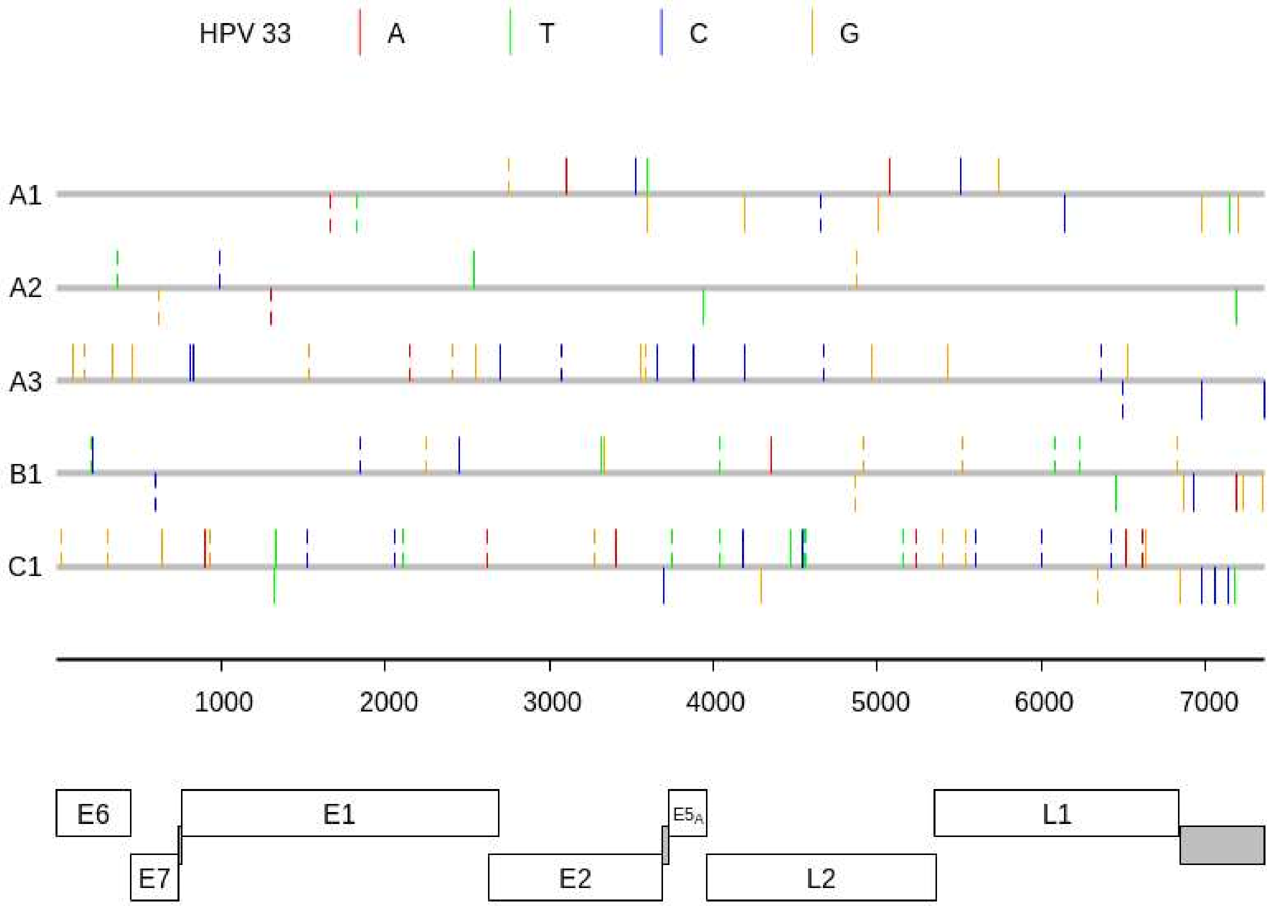

HPV 33
A
T
C
G
A1
A2
A3
B1
C1
1000
2000
3000
4000
5000
6000
7000
E6
E7
E1
E2
E5A
L2
L1

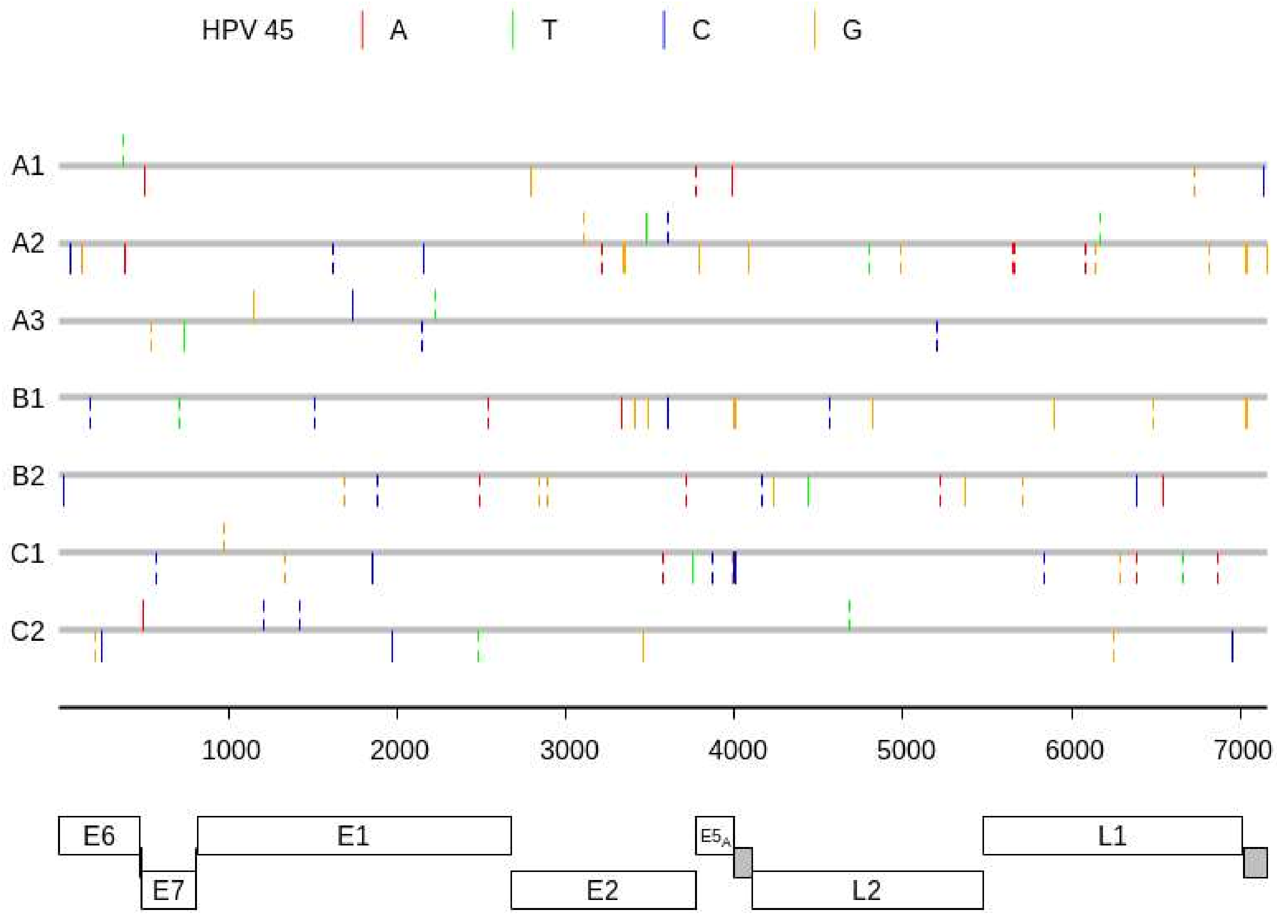
HPV 45
A
T
C
G
A1
A2
A3
B1
B2
C1
C2
1000
2000
3000
4000
5000
6000
7000
E6
E7
E1
E2
E5A
L2
L1

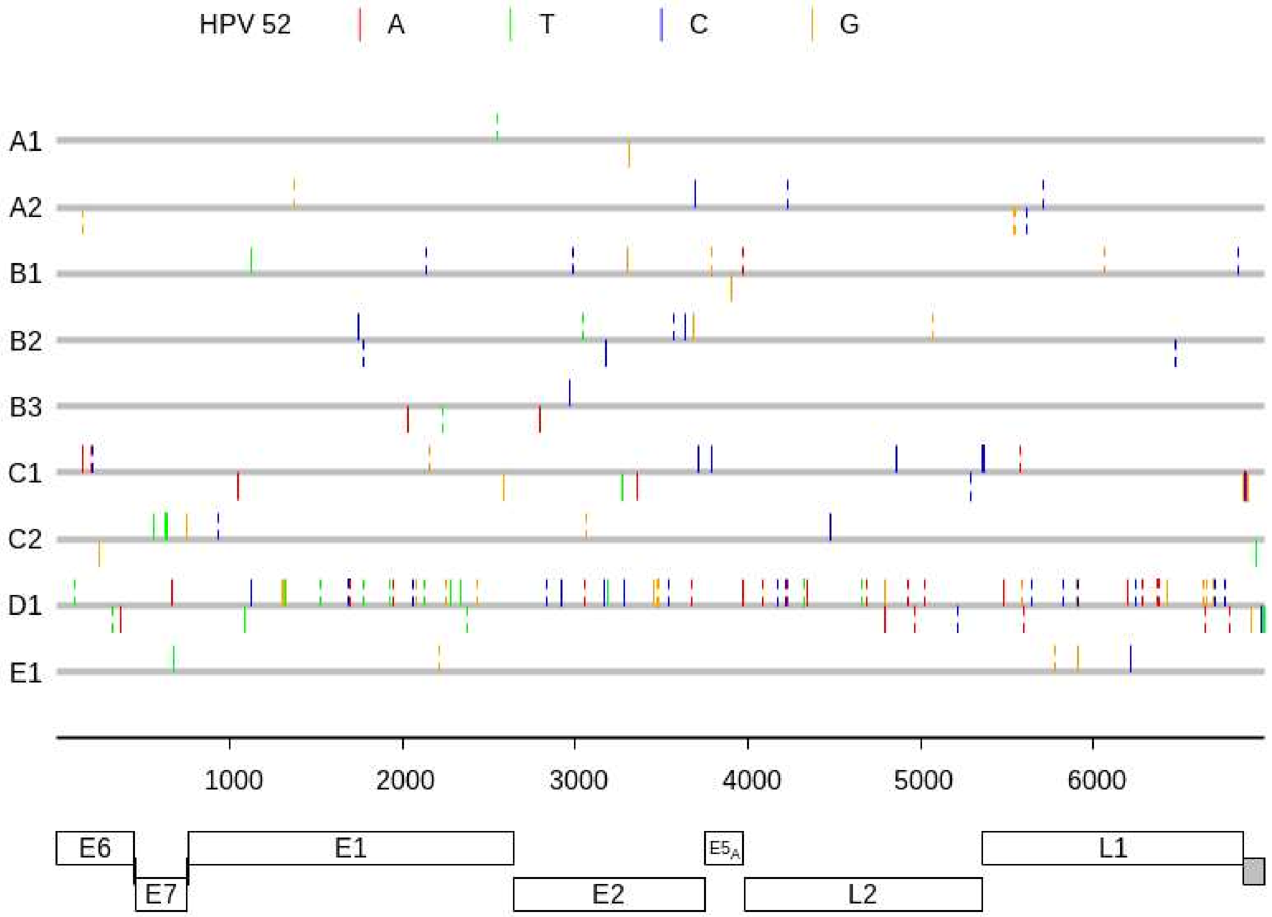
HPV 52
A
T
C
G
A1
A2
B1
B2
B3
C1
C2
D1
E1
1000
2000
3000
4000
5000
6000
E6
E7
E1
E2
E5A
L2
L1

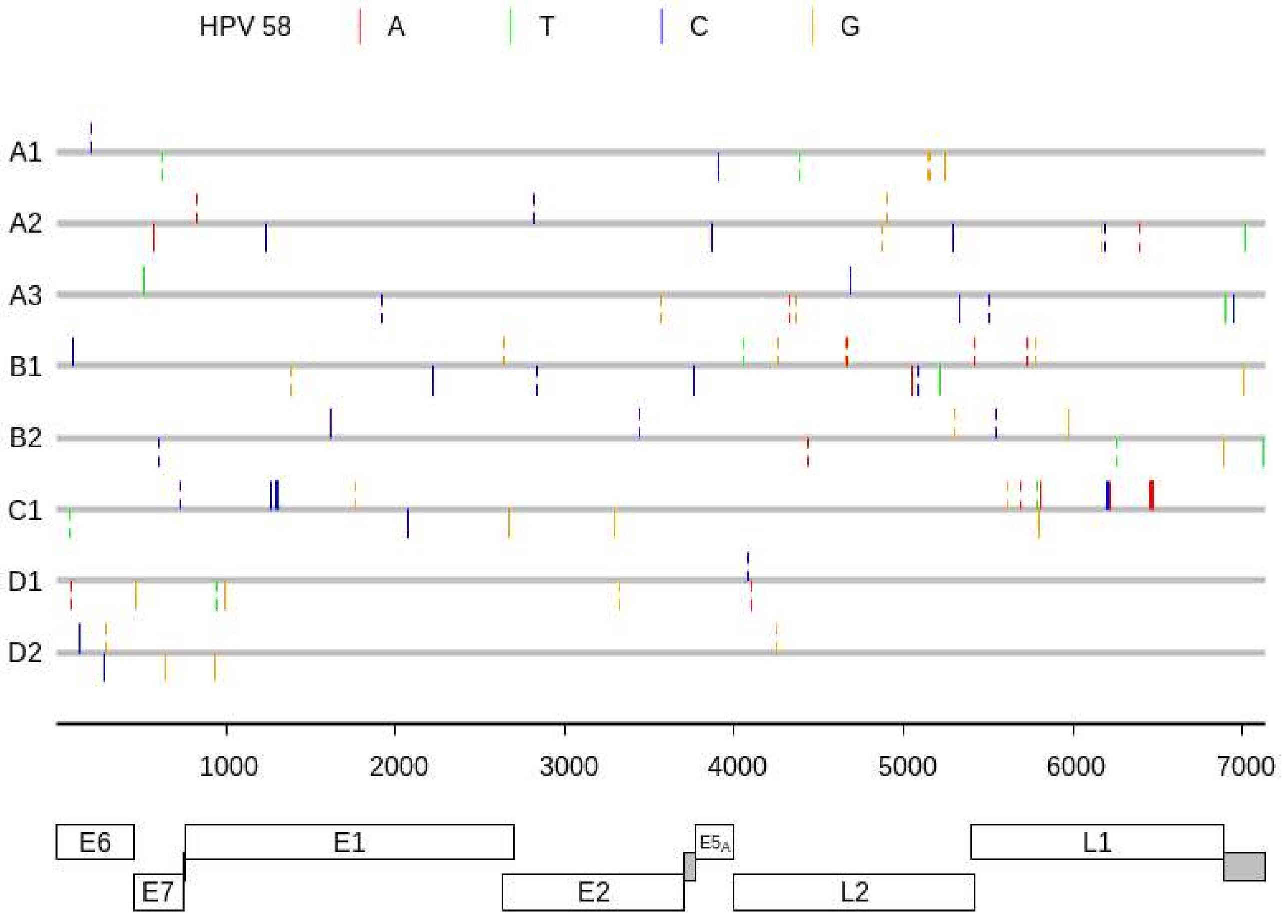

HPV 58
A
T
C
G
A1
A2
A3
B1
B2
C1
D1
D2
1000
2000
3000
4000
5000
6000
7000
E6
E7
E1
E2
E5A
L2
L1
